# Genomic history and selection in Roman and early medieval Britain

**DOI:** 10.64898/2026.04.28.721361

**Authors:** Marina Silva, Thomas Booth, Kyriaki Anastasiadou, Jesse McCabe, Christopher Barrington, Monica Kelly, Mia Williams, Yusuf Mohamedy, Mackenzie Masters, Alexandre Gilardet, Sarah Johnston, Frankie Tait, Pooja Swali, Jessica Peto, Isabelle Glocke, Sam Leggett, Lucy J. Koster, Helen Fewlass, Britta van Tiel, Emer Bealin-Kelly, Margaret R. Crawford, Oscar R. Aldred, Martyn G. Allen, Edward Biddulph, Christopher Booth, Ceridwen V. Boston, Adelle Bricking, Lisa Brown, Katherine Buckland, Ciara Clarke, Elizabeth Craig-Atkins, Neil G. W. Curtis, Alex Davie, Gabrielle Delbarre, Johan Du Preez, Laura H. Evis, Ceri Falys, Mark Gibson, David Gilbert, Katie A. Hemer, Andy Hood, Maya Hoole, Amal N. R. H. Kreisheh, Matthew G. Knight, Simon Mays, Keith Parfitt, Alexander J. E. Pryor, Rebecca Redfern, Kerry L. Sayle, Jo Seaman, Colin Shell, Sarah Stark, Megan C. Stoakley, Adelina C. Teoaca, Jess E. Thompson, Don Walker, Gaynor Western, Hugh Wilmott, Annsofie Witkin, Andrew Young, Sharon J. Clough, J. Alison Sheridan, Milena A. Grzybowska, Louise Loe, Malin R. Holst, Rick J. Schulting, Kate Britton, Gordon Noble, Ian Armit, Jo Buckberry, Ian Barnes, Selina Brace, Mark G. Thomas, Linus Girdland-Flink, Adrián Maldonado, Peter Heather, James C. Lee, Leo Speidel, Pontus Skoglund

**Affiliations:** Ancient Genomics Laboratory, The Francis Crick Institute, London, United Kingdom; UCL Institute of Archaeology, London, United Kingdom; Bioinformatics and Biostatistics, The Francis Crick Institute, London, United Kingdom; BioArCh, Department of Archaeology, University of York, York, United Kingdom; Archaeology, Environmental Changes & Geo-Chemistry Research Group, Vrije Universiteit Brussel, Brussels, Belgium; York Osteoarchaeology Ltd., York, United Kingdom; Centre for Palaeogenetics, Stockholm, Sweden; Department of Archaeology, University of Reading, Reading, United Kingdom; Research Department of Genetics, Evolution and Environment, University College London, London, United Kingdom; Department of Archaeology and History, University of Exeter, Exeter, United Kingdom; School of History, Classics & Archaeology, University of Edinburgh, Edinburgh, United Kingdom; Department of Archaeology, University of Aberdeen, Aberdeen, United Kingdom; University of Bristol, Bristol, United Kingdom; Genomics Science Technology Platform, The Francis Crick Institute, London, United Kingdom; Cambridge Archaeiological Unit, Department of Archaeology, University of Cambridge, Cambridge, United Kingdom; Oxford Archaeology Ltd, Oxford, United Kingdom; Wessex Archaeology, Salisbury, United Kingdom; Amgueddfa Cymru - Museum Wales, Cardiff, United Kingdom; Wiltshire Museum, Devizes, Wiltshire, United Kingdom; Eastbourne Borough Council, United Kingdom; AOC Archaeology Group, United Kingdom; School of History, Philosophy and Digital Humanities, University of Sheffield, Sheffield, United Kingdom; University Collections, University of Aberdeen, Aberdeen, United Kingdom; Down Farm Museum, Dorset and Bournemouth University, Faculty of Health, Environment and Medical Sciences, Bournemouth, United Kingdom; Thames Valley Archaeological Services Ltd, Reading, United Kingdom; Rubicon Archaeology Group Ltd, Cardiff, United Kingdom; Foundations Archaeology, Swindon, United Kingdom; Historic Environment Scotland, Edinburgh, United Kingdom; South West Heritage Trust, Taunton, United Kingdom; National Museums Scotland, Edinburgh, United Kingdom; Investigative Science, Historic England, Portsmouth, United Kingdom; Department of Archaeology, University of Southampton, Southampton, United Kingdom; Kent Archaeological Society, United Kingdom; London Museum, London, United Kingdom; School of History, Classics and Archaeology, Newcastle University, Newcastle, United Kingdom; SUERC, East Kilbride, Glasgow, United Kingdom; Friends of the Mint House, United Kingdom; McDonald Institute for Archaeological Research, University of Cambridge, Cambridge, United Kingdom; SLR Consulting Ltd., Carlisle, United Kingdom; Department of Archaeology, University of York, York, United Kingdom; Canterbury Archaeological Trust, Canterbury, United Kingdom; Department of Archaeology, Canterbury Christ Church University, Canterbury, United Kingdom; Museum of London Archaeology (MOLA), London, United Kingdom; Ossafreelance, United Kingdom; Avon Archaeology, United Kingdom; Cotswold Archaeology, Cirencester, United Kingdom; Archaeological Research Services Ltd., Bakewell, United Kingdom; School of Archaeology, University of Oxford, Oxford, United Kingdom; Archaeological and Forensic Sciences, University of Bradford, Bradford, United Kingdom; The Natural History Museum, London, United Kingdom; Department of History, King’s College London, London, United Kingdom; Genetic Mechanisms of Disease Laboratory, The Francis Crick Institute, London, United Kingdom; Institute for Liver and Digestive Health, University College London, London, United Kingdom; Center for Interdisciplinary Theoretical and Mathematical Sciences, RIKEN, Wako, Japan

## Abstract

Leading biomedical resources rely on genome variation in Britain^1–3^, but the historical processes that shaped present-day fine-scale diversity remain debated^4–13^. Here we sequenced 1039 ancient shotgun genomes from Britain (median 1.4-fold coverage), primarily dating to the first millennium CE. We imputed ∼660 million variants in the UK Biobank^14–16^ and employed genealogy-based ancestry reconstruction. We found an association between Iron Age consanguinity and matrilineal burial practices^17^, later disrupted following the Roman Conquest. Despite this societal impact, only 20% of Roman-period individuals carried detectable ancestry from outside Britain. In contrast, from the 6th century CE we detect widespread influx of ancestry in over 70% of individuals in southern’Anglo-Saxon’ Britain, with limited local admixture. We find previously underappreciated heterogeneity, with ancestries associated with Central and Southern Europe rising in prevalence from the 7th century CE. We demonstrate distinct Scandinavian-related ancestry in many Viking-associated contexts, but show that the population-level impact of the Viking Age in Britain was limited. Finally, we detect pre-medieval selection on variants linked with key immunity genes *TLR10-TLR1* and *IRF8*. These results identify population-level and selective processes that shape variation and disease risk in Britain today.

## Main Text

Studies of genetic variation in large medical biobanks represent a key avenue into understanding the link between genomes and human biology, including disease. However, genome-wide association studies linking variants to traits require an understanding of the genomic history that shaped genetic variation, as even subtle but unknown population structure can be an obstacle to understanding the genetic architecture of human biology^18,19^. Consequently, investigating the processes that shaped present-day population structure is not only of intrinsic historical interest but is also important for the accuracy of biomedical studies^20^.

Studies of present-day genetic variation in Britain have documented substantial fine-scale structure and hypothesised that it was largely shaped by pre-Roman movements, with less than half of present-day ancestry deriving from medieval expansions^4^. Although the Roman period in Britain has been largely undersampled, previous studies have highlighted the genetic impact of early medieval demographic processes in Europe^4,6,21–25^, with clear differentiation between medieval and earlier periods also found in Britain^6,8,9^. However, the timing and dynamics of movements of people across the North Sea, the circumstances by which Germanic languages were adopted in Britain and the later potential impact of population movements in the Viking Age are still not fully understood. This is due in part to the lack of reliable historical sources, particularly for the arrival of people identified as Saxons by Gildas, writing contemporaneously in the 5th and 6th centuries CE (the *Adventus Saxonum*)^26^. There are also challenges in discerning very closely-related ancestries, with advances in imputation approaches and high-resolution genomic methods tailored for aDNA having emerged only recently^21,27^.

The majority of aDNA data published from Britain is restricted to capture data based on ∼1.2 million previously ascertained loci, with only a limited number of shotgun whole-genomes from historic periods (**Extended Data Fig. 1**). This offers insufficient resolution to infer detailed historical population dynamics or how they influenced present-day genetic variation. Furthermore, studies of selection in West Eurasia over the past 10,000 years^28^ (using both SNP data from 1.2M loci and whole-genome sequences)^7,28–31^ did not focus on narrow time periods and may have been affected by ancestry changes.

### Dense time transect of whole genomes from Britain

We sequenced 1039 genomes from Britain (median 1.4-fold coverage), spanning from 2550 BCE, in the Bronze Age, to 1150 CE, after the Norman conquest. Our main focus was the 1^st^ millennium CE, particularly the Roman and early medieval periods, which are recognised as times of societal change and increased migration, as well as the immediately preceding and succeeding transitional periods (*n*=834, between 50 BCE and 1150 CE) (**Fig. 1a**). We also included 200 Bronze Age and Iron Age genomes to generate a comparative whole-genome dataset (**Extended Data Fig. 1; Extended Data Table 1;** five additional individuals were later dated to more recent periods).

**Fig. 1:**
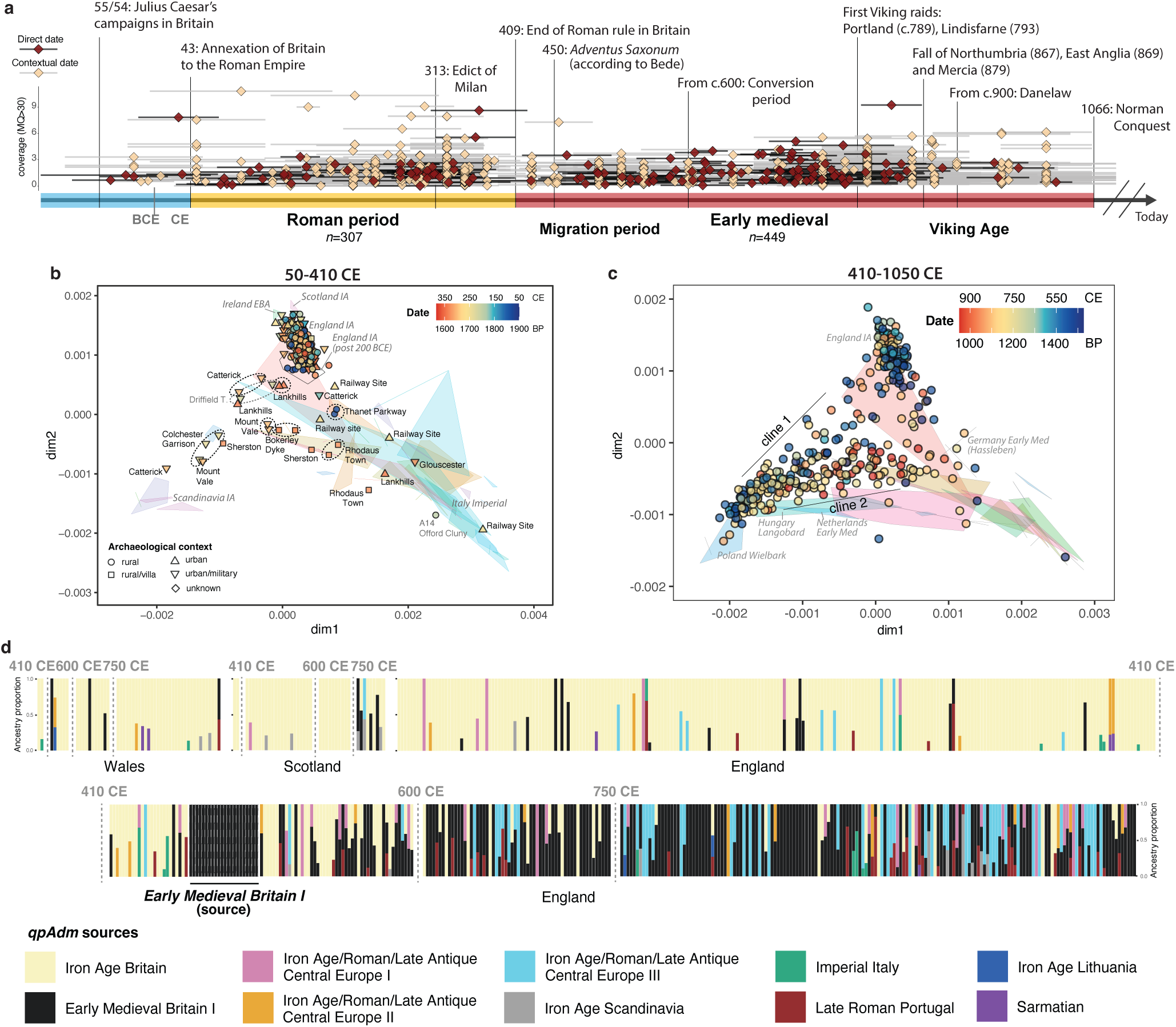
Ancestry changes in Britain in the 1st millennium CE. **a)** Temporal distribution of newly-sequenced samples, with average coverage (after restricting to sequencing reads with mapping quality (MQ) above 30) and simplified timeline of main historical events in Britain during the 1st millennium CE. Directly radiocarbon dated individuals in dark red. Error bars denote 1 standard error. Individuals dating to Bronze and Iron Ages, as well as post-dating 1066 CE not shown in the timeline (**Extended Data Table 1; Extended Data** Fig. 1a). **b)** MDS based on *Twigstats*-boosted pairwise outgroup-*f*3s (*t*=1000 cutoff) focussed on the Roman period. Polygons represent individuals from continental Europe, Britain and Ireland dating to the Bronze and Iron Ages and the Roman period. **c)** MDS based on *Twigstats*-boosted pairwise outgroup-*f*3s (*t*=1000 cutoff) focussed on the early medieval period. Polygons represent individuals from continental Europe from Roman and early medieval periods. In b) and c) data points are coloured according to the average date in BP of each individual. **d)** Twigstats-boosted *qpAdm* models for individuals from Britain dating to the 1st millennium CE. Both newly sequenced and previously published individuals are included and are plotted according to their average date (**Extended Data Table 1**); note that the individuals included in the *Early Medieval Britain I* source (all dated to the 6th century CE) are also plotted. For site labels and specific individuals highlighted in the main text see **Supplementary** Fig. 1. Only individuals with accepted *qpAdm* models are shown (*p*-value>0.01; **Methods**). Full *qpAdm* results in **Extended Data Table 8**.

We used a single-stranded DNA library preparation protocol^32^ and computationally removed post-mortem damage in a strand-dependent manner prior to imputation (**Methods**), removing damage-induced biases and improving imputation accuracy (**Extended Data Fig. 1**). This approach also allows accurate imputation of postmortem damage in CpG contexts, which account for over a quarter of common human SNPs but, due to methylation, are uncorrected by approaches that rely on enzymatic treatment targeting uracil residues^33^. We imputed ∼660 million variants using the UK Biobank reference panel^15,16^ and inferred genealogies^34^ from a total set of 1722 ancient genomes (**Methods**) with a coverage of at least 0.5-fold, and restricted genealogies to recently-shared ancestry with Twigstats^21^.

Although previous DNA sampling from Roman Britain has been relatively small-scale and regionally or context specific^9,35,36^, our dataset bridges this gap. Whilst we find broad continuity with Iron Age Britain (**Fig. 1b,d; Supplementary Fig. 1**), by contrast, we detect a large-scale arrival of ancestries, which we associate with Germanic-speaking groups, in the early medieval period, although with higher heterogeneity than previously described^6,8,9^ (**Fig. 1c,d**; **Supplementary Fig. 1**). We investigate the temporal and geographical patterns of ancestry and relatedness throughout the 1st millennium CE in detail.

### Ancestry and relatedness in Iron Age and Roman Britain

Southeastern Britain fell within the Roman sphere of influence prior to Julius Caesar’s failed attempts to invade in 55 and 54 BCE, but the Roman annexation of south and east Britain was only achieved later, as a result of campaigns starting in 43 CE led by the Emperor Claudius, with Roman rule lasting until ∼410 CE^37^ (**Fig. 1a**). In other parts of Europe, the Roman period witnessed political and societal changes resulting in an intensification of human mobility, as evidenced by aDNA and isotopic studies^9,38–42^, facilitated by the development of road and maritime networks associated with military campaigns and trade. Archaeological evidence indicates that southern Britain was integrated within these networks, for example through the establishment of urbanism, villas, varied industries, Romanised funerary practices, and the presence of imported goods from across the Roman Empire^37,43^. Later in the Roman period, funerary traditions and religious practices changed with the wider adoption of Christianity following the Edicts of Nicomedia and Milan in 311 and 313 CE (**Fig. 1a**)^44^. Although the long-term transformative economic, political and cultural effects of incorporation into the Roman system are clear both in urban and rural regions^12^, including in some sites beyond the frontier north of the Hadrian and Antonine Walls^45^, prior to this study genomic data was available from only 10 archaeological sites in Britain dating to this ∼350-year period^9,10,35,36^ and thus the genetic impact on local communities in different regions and contexts remained unclear^46–48^.

The preceding Iron Age in Britain (800 BCE–43 CE) is traditionally regarded as a period of increased insularity and regionalism, despite some evidence from both archaeology and genetics for cross-channel contacts between southeast Britain and, possibly, confederations from Gaul (*e.g.* the Belgae)^17,49^. However, it has been unclear how Iron Age communities were initially affected by Roman rule.

Previous work identified evidence for pervasive matrilocality in several Iron Age sites displaying reduced mitochondrial DNA (mtDNA) diversity^17^. Here we identify a similar pattern in one additional Iron Age site: Webbington Farm (Hinkley, Somerset) (**Fig. 2a**). Overall we observe higher within-site Identity-By-Descent (IBD) in sites where we also detect low mitochondrial haplogroup diversity, consistent with matrilineal burial practices (**Fig. 2b; Extended Data Fig. 2; Supplementary Fig. 2; Extended Data Table 2; Methods**). Making use of the larger sample sizes in Yorkshire, we compared patterns of IBD sharing and mtDNA haplogroups within and between Wetwang Slack (*n*=79), and the neighbouring ‘Arras’ cemeteries of Pocklington (*n*=30) and East Coast Pipeline (*n*=7)^10^, all <25 km from each other. All three sites display low mtDNA diversity, but with different haplogroups dominating and little overlap across sites, despite high IBD-sharing between the sites, consistent with an extreme scenario of matrilineal burial practices at very small geographical scales (**Extended Data Fig. 2**). Relatives detected between any two sites (all above 3rd degree) did not share a maternal lineage (an additional pair of distant related individuals across Wetwang Slack and Nunburnholme Wold, within 13 km, also carried different mtDNA haplogroups). These results could be explained by women tending to reside near their maternal relatives (assuming burial location would reflect residence), resulting in little contact between sites along the female line of descent, or alternatively a scenario in which women left their place of birth but after death their bodies and those of their descendants’ were returned to be buried near their maternal relatives. However, burial practices alone would not be expected to result in increased IBD-sharing (**Fig. 2b**). While a recent study showed that this kinship system was maintained at Wetwang Slack for multiple generations within several large genealogies (Olalde *et al*. 2026 *in prep*), within-site sample sizes are too small to formally test this hypothesis in other regions.

**Fig. 2:**
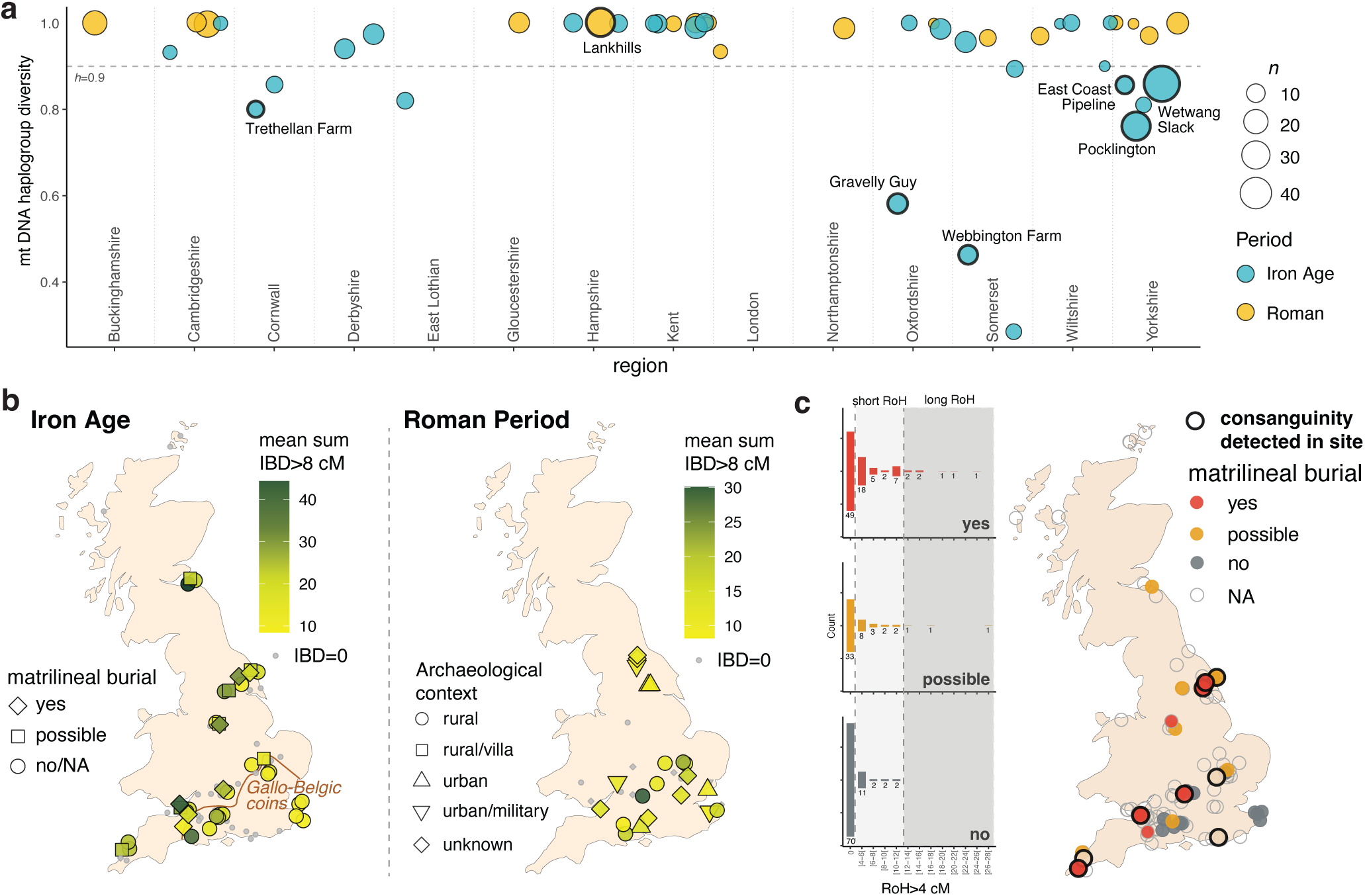
Iron Age regionalism and disruption of kinship practices in Roman times. **a)** Haplogroup diversity (h) within Iron Age and Roman-period sites with n≥5, after filtering out related individuals up to 3rd degree. Sites are grouped by region and coloured according to period (h calculated using a combined dataset comprising newly and previously reported individuals; **Extended Data Table 2**). Sites with at least one consanguineous individual are labelled. **b)** Mean pairwise IBD-sharing within sites in the Iron Age (left) and Roman period (right). Sites are coloured according to within-site pairwise IBD-sharing >8 cM, with sites with *n*=1 or with pairwise IBD-sharing=0 shown in grey. **c)** Frequency of ROHs (left) in sites with different signals of matrilineal burial practice (yes, no, possible; **Methods**). Map of Iron Age sites (right), coloured according to the signal observed on mitochondrial haplogroup signals of matrilineal burial practice. Sites with at least one consanguineous individual indicated with thick black outline.

Individuals from cemeteries where we detect low mitochondrial diversity displayed more short runs of homozygosity (ROH 4-8 cM) (Kruskal-Wallis test: *p*=7.558e-4), indicating lower effective population size (**Fig. 2c**; **Extended Data Table 3 and 4**). We also detect a statistical association between sites with a low mitochondrial diversity signal and consanguineous practices (**Fig. 2c**; **Extended Data Table 3 and 4;** Fisher’s Exact Test: *p*-values ranging from 0.003 to 0.021), with individuals who are the offspring of consanguineous unions defined as carrying >50 cM in total of runs of homozygosity (ROHs) longer than 12 cM (likely representing the offspring of first cousins or more distantly-related parents)^10^ (**Extended Data Table 5**). These close-kin unions, although probably sporadic (**Extended Data Fig. 2**), could either be an outcome of limited population sizes (possibly linked to a system of social stratification involving a section of society who largely had children among themselves) or result from practices used to reinforce specific kinship systems. An association between matrilineality and endogamy has been previously described in both present-day and ancient populations^50,51^. Remarkably, two of three individuals resulting from consanguineous unions identified in our Roman-period dataset were found at sites in Somerset (C13400, Hole Ground, Wookey Hole) and Dorset (C11612, Woodcuts), both regions where we detect low intra-site mitochondrial diversity in the preceding Iron Age (**Extended Data Table 2 and 3**), suggesting the possibility of occasional continuity of social practices. However, neither of these skeletons has been directly dated, and given the wider archaeological context at both sites, it is possible that these date to the Iron Age instead. One additional Roman-period individual showing genetic signals of consanguinity has been reported from Dorset^17^.

We find that the organisation of cemeteries in the Roman period did not reflect a straightforward unilateral descent system, suggesting that burial practices inferred in some Iron Age^17^ sites were modified or abandoned during the Roman period (**Fig. 2a; Supplementary Fig. 3**). The disruption of previous kinship practices could be explained by the establishment of Roman political structures, typically patriarchal in nature, overturning specific customs and practices in a newly annexed province. We note however that we do not have good sampling from Roman-period Cornwall, Dorset or from rural sites in Yorkshire – all regions with a strong matrilineal signal in the Iron Age – to formally assess continuity in funerary practices within these regions.

The patterns of IBD-sharing and relatedness within and between sites offer an additional perspective on mobility within Britain during the Roman period. We identified seven pairs of distantly-related individuals across different Roman-period sites, five of which are within 50 km of each other (**Extended Data Table 6**). Despite the small sample size, we detect a clear pattern: all pairs of relatives found between rural sites were within a <15 km radius, whereas all relatives found between sites >40 km apart involved at least one urban or military cemetery, suggesting increased scales of movement and connection associated with urban or military sites.

Mean pairwise IBD sharing within sites in the Roman period was overall lower than in the preceding Iron Age, evidencing disruption of previous regional social structure. IBD sharing was higher within rural than urban or military-associated cemeteries (**Fig. 2b; Extended Data Fig. 3**). Similarly, we detect a low proportion of close relatives (*i*.*e*. up to third-degree; **Extended Data Table 7**) in cemeteries serving large urban settlements (*e*.*g*. Lankhills, Railway Site), although we caution our current DNA sampling might be less representative of whole large cemeteries. We find a similar general trend in military-associated sites, although we also identified a father-son pair in Gloucester, a *colonia* for legionary veterans and their families dating to 250-410 CE, both with ancestry related to Iron Age Britain, consistent with accounts of military recruitment from the local population^52^. In contrast, despite the smaller sample size, we identified genetically closely-related communities in smaller cemeteries in rural locations, such as Childrey Warren in Oxfordshire, with high mean intra-site pairwise IBD-sharing (∼30 cM; **Fig. 2**) and six of eight individuals in close genetic relationships (3rd degree or less) (**Extended Data Fig. 3b**), although notably, patterns of relatives resembled a bilateral rather than matrilineal-based burial system.

This change in relatedness patterns occurred against a background of widespread continuity in ancestry from the Iron Age into the Roman period: 71% of 259 individuals dated to the Roman period are modelled with Twigstats as tracing 100% of their ancestry to Iron Age Britain (100% *Iron Age Britain pre-200 BCE*; *p*>0.01) (**Fig. 1d; Extended Data Table 8**). This rises to 80% of individuals when including a second source comprising Iron Age individuals from southeast Britain post-dating 200 BCE whose ancestry was shifted towards nearby regions of continental Europe (*Iron Age Britain post-200 BCE*) (**Extended Data Fig. 4a; Extended Data Table 9**). This lack of substantial influx of ancestry in the Roman period is also supported by a relative absence of long IBD segments (>25 cM) detected between individuals in Britain and Europe (**Supplementary Fig. 4**).

In contrast, ∼20% of individuals carried disparate ancestries, consistent with individual transient mobility, similar to what has been observed in other Roman provinces^42^. These ancestry outliers date mostly to the 3rd and 4th centuries CE (but we caution that this could reflect a relative lack of inhumations and thus fewer opportunities for aDNA sequencing in the earlier Roman period^53^; **Extended Data Fig. 4c**) and carried an array of ancestries, denoting contributions from northern and central European, Mediterranean and Steppe-related sources (**Fig. 1d**). These individuals were primarily found in urban centres and/or in contexts associated with the military (Lankhills in *Venta Belgarum*, present-day Winchester; Railway site, Mount Vale, and the previously published Driffield Terrace^9^ in *Eboracum*, present-day York; Bainesse Farm in Catterick; Colchester Garrison in *Camulodunum*; Rhodaus Town in *Durovernum Cantiacorum*, present-day Canterbury; and Gloucester, *Glevum*), and at villa sites (Sherston, in Wiltshire, and Bokerley Dyke, in Dorset), adding to previous observations from Britain^9,35^ (**Fig. 1b**; **Extended Data Fig. 4c; Supplementary Fig. 1**). Some of these sites, such as Catterick, Lankhills and Railway site, displayed high heterogeneity in ancestries, in line with the Roman policy of garrisoning the frontiers by non-local soldiers^54^, and together with isotopic data^55,56^, document long-distance connections between Britain and different parts of the Roman Empire, and potentially beyond. However, ancestry outliers were also detected in rural sites at Thanet Parkway (Kent), Childrey Warren (Oxfordshire) and A14-Offord Cluny^36^ (Cambridgeshire), suggesting this long-distance mobility did have a sporadic, albeit seemingly minor, influence on rural areas (**Fig. 1b**; **Extended Data Fig. 3c**).

### Early medieval ancestry transformation

Decades of political instability in the wider Roman Empire eventually led to the end of Roman administration in Britain (**Fig. 1a**). The ensuing Migration Period (∼400-600 CE), saw the fall of the Western Roman Empire, and was characterised by movements of groups across Europe, particularly of Germanic-speakers^57^. Germanic languages likely began being spoken persistently in Britain at this time, eventually giving rise to the English language. Funerary practices, architecture and material culture show clear links with continental regions across the North Sea spanning present-day France, Frisia, Jutland, Sweden and parts of northern Germany^58^, and while both genetic^6,8,9^ and isotopic^59^ studies have inferred large-scale movement of people into Britain, the nature and timings of these contacts are still debated.

A recent genetic study has shown that this period was associated with an influx of ancestry maximised in continental northwestern Europe^8,9^, as well as ancestry modelled with a source related to Iron Age France; the latter reported as restricted to southern Britain^6^. High-resolution Twigstats-based MDS analysis corroborates this result, showing the bulk of individuals with a mean date (either radiocarbon or contextual) post-dating ∼410 CE clustering away from the main preceding Roman-period individuals, indicating a major influx of ancestries that were previously rare or absent in Britain. However, we detect substantial heterogeneity among these arriving ancestries, visible as two major clines, with a striking temporal pattern: cline 1, spanning from an Iron Age Scandinavia cluster to the Iron Age and Roman Britain cluster, and cline 2, with mostly later individuals, extending from the end of cline 1 to a Roman and Late Antique Central European clusters (**Fig. 1c**).

There is considerable debate on the extent to which the archaeological changes of the 5th century CE can be attributed to large-scale migrations^6,13,47,58–61^. Although we confirm that continental-related ancestries were sporadically present in Britain during the Roman period, they were mostly restricted to urban, military or elite sites (**Fig. 1b**). The low proportion of Roman-period individuals carrying such ancestries is probably insufficient to explain the magnitude of ancestry transformation observed in the medieval period. It is only after the end of Roman rule that these ancestries became widespread across large parts of Britain (**Fig. 1d**).

We applied the *qpAdm* framework implemented by Speidel et al^21^ with a modified set of sources (**Methods**) to model ancestry for each individual from Britain dating to the 1st millennium CE (**Extended Data Table 8**). We tested all combinations of one, two or three Iron Age, Roman and/or Late Antique putative ancestry sources from Britain and continental Europe. We added a new source population – *Early Medieval Britain I* – comprising the earliest individuals in Britain clustering at the end of cline 1 (**Fig. 3**), all dating to the 6th century CE, who also showed similarly distinct ancestry profiles in a genetic clustering analysis (*K*=6) (**Supplementary Fig. 5**; **Methods**). This group of individuals includes the Marlow Warlord burial (C10756), a richly-furnished grave found near Bisham, in Berkshire, and Sk 101 (C11210) from the Anglo-Saxon cemetery at Three Kings, in Cambridgeshire, who we identify as a distant (4th-6th degree) genetic relative of MDM007^6^, found in the site of Midlum, in Frisia (present-day Netherlands) – both showing clear archaeological or genetic links with the North Sea shore. We thus emphasise that despite its label, this ancestry probably traces back to continental Europe, and did not originate in Britain. The ancestry maximised in this group of individuals is differentiated from that maximised in Iron Age Scandinavia at higher *K* values (**Supplementary Fig. 5**). While it might have been expected that early medieval individuals in Britain, possibly Saxons, would derive at least some ancestry from Iron Age Scandinavia^21^, we show that they are clearly distinct in the Twigstats-boosted MDS and *Iron Age Scandinavia* is rarely accepted as a source in our *qpAdm* modelling (**Fig. 3; Supplementary Fig. 6**).

**Fig. 3:**
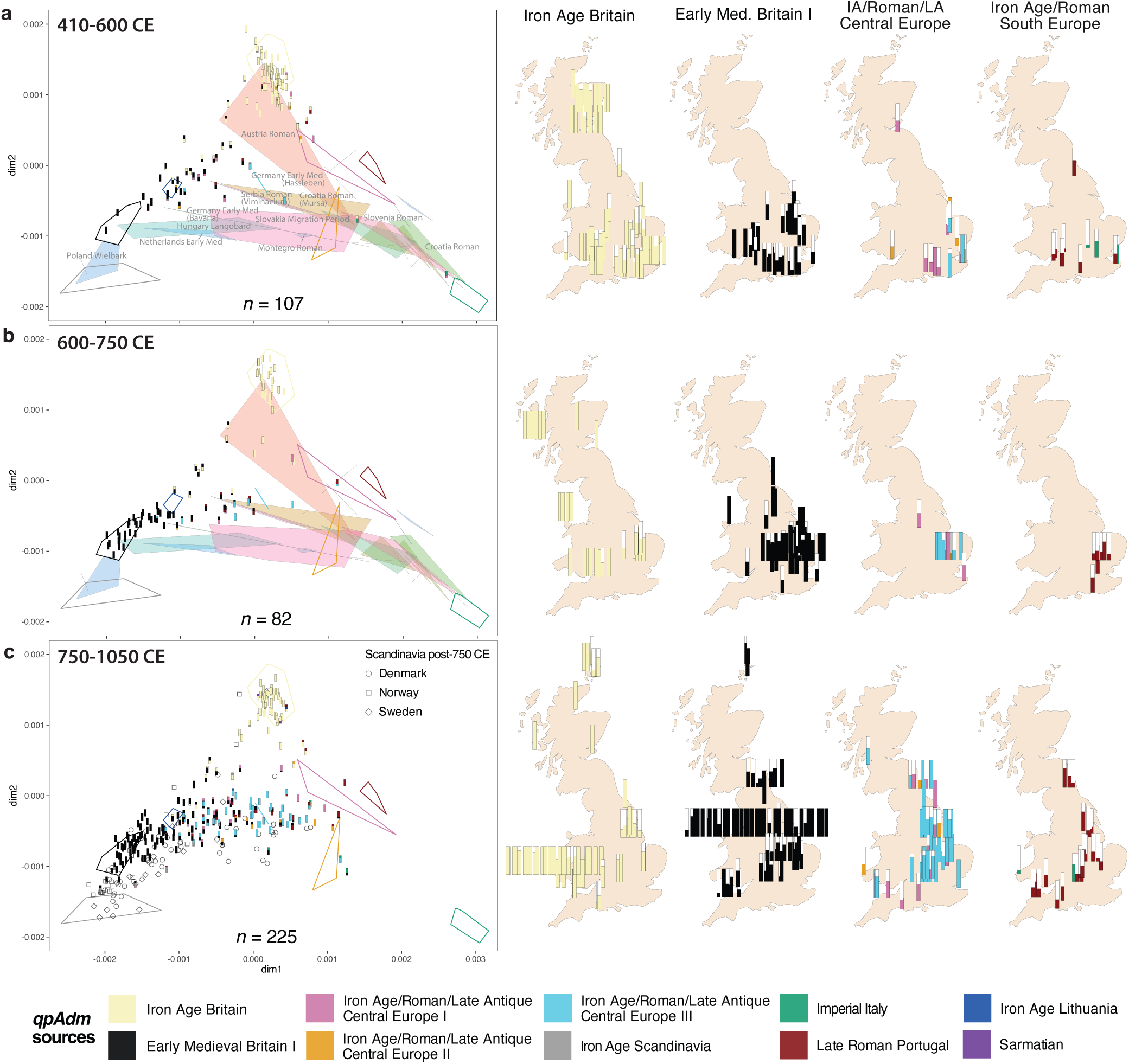
Influx of ancestries in the early medieval period: **a)** 410-600 CE, **b)** 600-750 CE, **c)** 750-1066 CE. MDS based on Twigstats-boosted pairwise outgroup-*f*3s (*t*=1000 cutoff), stratified by time. Individuals from Britain (both newly sequenced and published) are used as targets and represented by their qpAdm modelling proportions (as in Fig. 2; showing only plausible and non-rejected models, i.e. *p*-value>0.01 and *ancestry proportion±SE* between 0 and 1). Number of targets (*n*) for each time period indicated on each MDS. Polygons represent source populations included in the *qpAdm* models (*Sarmatian* not included in the MDS) and previously published continental populations dating to the Roman and early medieval periods. Maps show the proportion of ancestry derived from each tested source for each individual (proportions of *Iron Age Scandinavia*, *Iron Age Lithuania* and *Sarmatian* not sources shown; see Supplementary Fig. 6). Model selection detailed in **Methods**. All *qpAdm* results in **Extended Data Table 8**.

In the 5th and 6th centuries CE, a substantial proportion of individuals in our dataset carried *Early Medieval Britain I* ancestry. However, only a low proportion of individuals displayed admixed ancestry from *Iron Age Britain* and *Early Medieval Britain I* (**Fig. 1d**; **Fig. 3a**), consistent with previous models of the early Anglo-Saxon transition^59,62^. Although we detect ancestries relating to Central and Southern Europe as early as the 5th and 6th centuries CE (mostly in sites near the southern and eastern coasts), in this period the majority of individuals sampled from what is now England could be modelled as deriving all or most of their ancestry from the *Early Medieval Britain I* source (**Fig. 3a**).

However, from the 8th-10th centuries CE *Early Medieval Britain I* ancestry became less prevalent, with many individuals instead carrying ancestries associated with Central, and to a lesser extent Southern Europe, a large proportion of whom can be modelled as deriving 100% of their ancestry from such sources (**Fig. 1d**; **Fig. 3c**). Isotopes also point to changing migration patterns starting from the 7th century CE^59,63^, but our sampling size in the 7th century is substantially lower compared to the later centuries and we still detect a higher prevalence of *Early Medieval Britain I* ancestry between 600-750 CE (**Fig. 3b**). The individuals in our dataset that postdate the Norman conquest in 1066 CE (*n*=69 with accepted Twigstats models), follow the same trend. We see no apparent impact on ancestry following the Norman conquest in this small sample size (**Supplementary Fig. 6**), consistent with a scenario of elite transfer^64^, with the caveat that the majority of these individuals come from a single site (Leicester Waterside), within the region in eastern Britain under Danish Viking control in the late 9th–11th centuries CE (the Danelaw).

The most common best-fitting Central European source (*Iron Age/Roman/Late Antique Central Europe III*) comprises individuals from a 4th century cemetery in Sarrebourg, present-day France, potentially linked to Alemannic Kingdom^42^. However, whilst this is the closest proximate source available it might not equate to the true source population (although it aligns with recent isotopic findings of increased, possibly female-mediated, movement from the Rhineland regions in the 7th and 8th centuries CE^63^). The exact genetic sources of these Central Europe-related ancestries are difficult to pinpoint with the available data. Considering current gaps in published whole-genome shotgun datasets, missing sources could include people living in regions of present-day northwestern Germany ∼500-600 CE. Alternatively, this signal could represent the fusion of ancestries during the 4th-6th centuries CE elsewhere in continental Europe. These results mirror recent findings of Central Europe-associated ancestries expanding into southern Scandinavia prior to the Viking Age, ∼800 CE^21^. Furthermore, an influx of ancestries from further south was also inferred in cemeteries in present-day Germany at the onset of the Migration Period^22,65^ and in multiple sites along the North Sea coast^6^. It is worth noting that the temporality, geographical spread and frequency of this signal in our dataset might be profoundly affected by funerary practices that render some individuals archaeologically invisible (*e*.*g*. depositions in rivers^60^) or by the widespread practice of cremation in both eastern Britain and continental northern Europe^13,57^.

We must therefore consider different, although not mutually exclusive, scenarios to explain our genetic results. One possibility is a roughly simultaneous 5th-6th century arrival of people carrying differentiated ancestries (*Early Medieval Britain I* and diverse Central Europe-associated), with groups predominantly carrying Central Europe-related ancestries primarily practicing cremation and only becoming more prominent in the genetic dataset after adopting inhumation, possibly following conversion to Christianity in the 7th century CE^66^. A second scenario is a later arrival of new groups carrying Central Europe-related ancestries. Such an event could perhaps have been obscured by historians such as Bede (writing in the 8th century CE), whose work framed the arrival of putative different groups (Saxons, Angles, and Jutes) as largely simultaneous in the 5th and 6th centuries CE, in contrast to earlier accounts by Gildas that do not mention any tribal identities other than the Saxons (Bede (*Historia Ecclesiastica*): I.15 (pp. 26-28); Gildas (*De Excidio*): I.23 (pp. 17-18))^26,61^. A third possibility is continued migration across the North Sea gradually introducing differentiated ancestries (in agreement with recent isotopic results^59^), reflecting ancestry changes in northwestern continental source groups, possibly as a result of earlier desertion of Germanic settlements in the same region^67^. Our genetic results add to the uncertainty and discussions around the accuracy of Bede’s accounts of the origins of Anglo-Saxon societies, including the ways in which these 7th-8th centuries CE societies thought about their origins, identities and connections, when Bede was writing.

### Low impact of continental ancestries in the north and west

From the 8th century CE, most of present-day England, and parts of what are now southern Scotland and the Welsh Marches were under the control of culturally Anglo-Saxon elites, with the establishment of a multitude of kingdoms whose centres of power, borders and alliances shifted through the centuries. Outside of these regions, historical toponyms, grave goods and funerary practices do not show clear and widespread links to Germanic-speaking regions in continental Europe^37,68^. To the west, Brittonic-speaking kingdoms remained linguistically and archaeologically distinct until the later medieval period^69^. North of the river Humber, the Old English language spread to the Firth of Forth and as far as modern Ayrshire in the west by the 8th century CE following the steady expansion of the Northumbrian kingdom, though parts of the west remained Brittonic-speaking through to the end of the millennium. Gaelic had been spoken in the west at least as early as the 6th century CE but possibly centuries earlier, and gradually spread eastward into the Pictish territories north of the Forth and eventually into formerly British regions^70^. Funerary practices remained highly regionalised across northern and western Britain, though these regions shared certain essential aspects, mainly a rejection of grave furnishings and the use of east-facing inhumation burial in cists, coffins or low mounds.

In the north of Britain (broadly corresponding to present-day Scotland), a region which saw limited Roman intervention and seemingly much less (large-scale) population movement in the Migration Period, the majority of individuals can indeed be modelled solely with Iron Age Britain (**Fig. 1d**; **Fig. 3**). Notable exceptions in the 5th-6th centuries CE, carrying disparate ancestries in combination with ancestry related to Iron Age Britain, comprise C10189 from Thornybank (southeastern Scotland), buried in a traditional long cist cemetery, who carried ancestry related to a Central Europe source, and two individuals from the Pictish-associated long cist cemetery of Lochhead Quarry (discussed below). Later individuals with disparate ancestry profiles were probably connected to Viking-related movements (**Fig. 4b; Supplementary Fig. 1**).

**Fig. 4:**
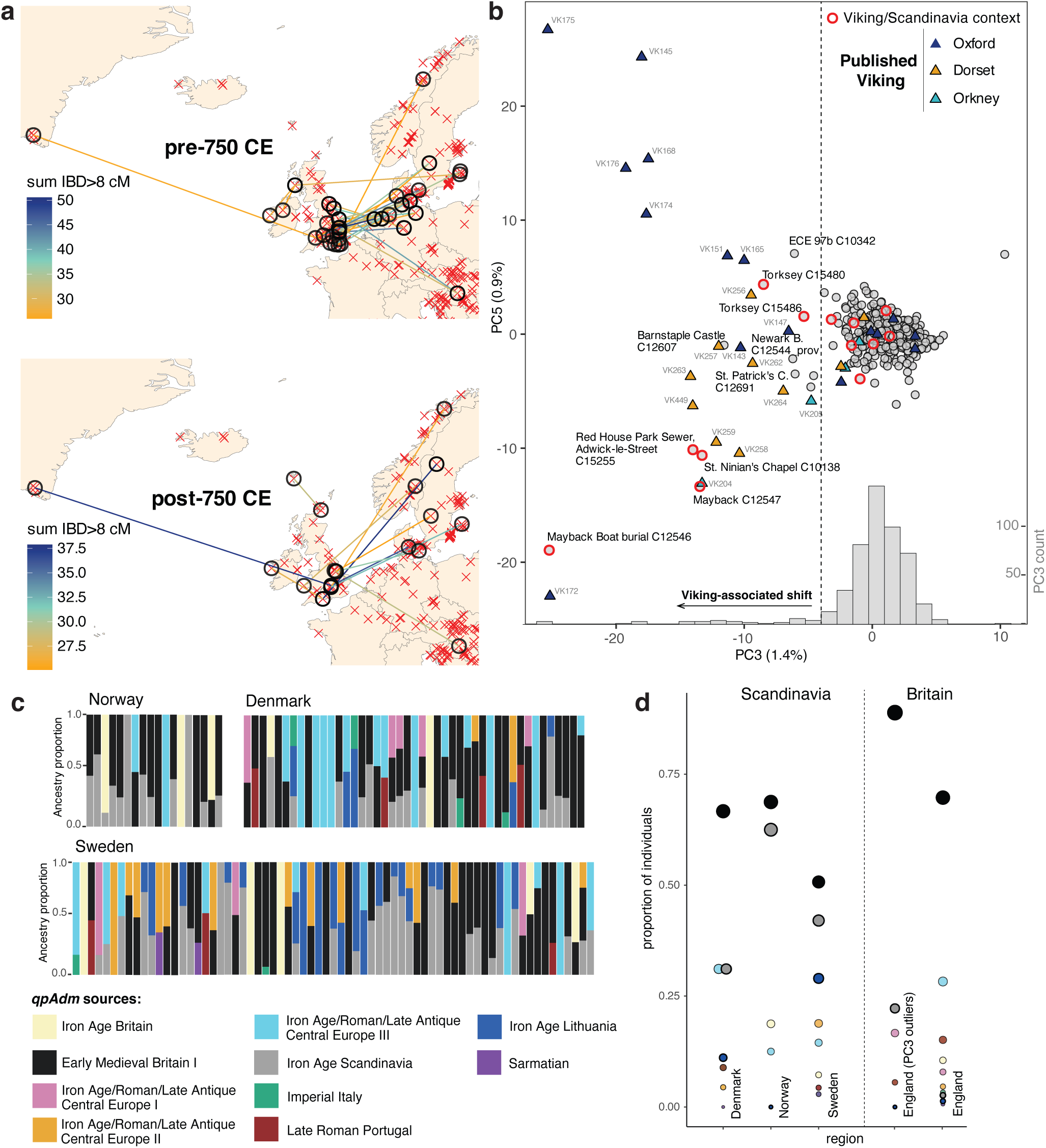
Chromosome segment sharing reveals early medieval and Viking Age recent ancestry links. **a)** IBD-sharing between medieval individuals in Britain (pre-and post-750 CE; 410-750 CE and 750-105 CE) and outside of Britain (dating to the Iron Age to the early medieval period). Sites with one or more individuals sharing at least one IBD segment >25 cM with another individual in a different site outside of Britain are represented by circles, and both sites are connected and coloured according to total IBD sharing (>8 cM). Remaining sites included in the analyses are represented by a single red ‘x’. Dataset includes shotgun and capture genomes (**Extended Data Table 6**). Note that these maps do not show IBD-sharing within Britain or between pairs that do not include one individual from Britain. **b)** PCA on Chromopainter-like genome-wide copying proportions, summarised using non-negative least squares (NNLS). Each target individual genome (from Britain, dating to the early medieval period; 410-1050 CE; grey data points) was painted against a palette comprising older genomes from Britain plus older and contemporary genomes from continental Europe (**Extended Data Table 13**). Individuals from previously published putative Viking contexts from Britain^23^ represented by triangles, coloured according to location. Newly-sequenced individuals from putative Viking contexts or sites with archaeological links to Scandinavia are highlighted with a red outline. Inset histogram of PC3 coordinates (y-axis shown on the right, in grey); vertical dashed line separates PC3 outliers (calculated as *Q1 - 1.5 x IQR*), which we label as the Viking-associated shift. **c)** Twigstats-boosted *qpAdm* models for individuals from Scandinavia dating to post-750 CE. Models for individuals from Britain are shown in Fig. 1d. **d)** Proportion of individuals in Scandinavia and Britain dating to the period 750-1050 CE carrying >10% ancestry from a given source. Size of data points represents the proportion of individuals within a certain region. *Early Medieval Britain I*, *Scandinavia Iron Age* and *Lithuania Iron Age* highlighted with a darker outline.

Building on the genomic analyses by Morez et al.^71^ we extend on existing Pictish aDNA data with the largest site-level genomic series reported to date, comprising twelve individuals from Lochhead Quarry (*n*=10 included in Twigstats analyses) that derive from a series of temporally well-dated contexts^72^. The majority of the Lochhead Quarry individuals could be modelled with only *Iron Age Britain* ancestry, consistent with earlier findings^71^, but for two individuals (C13045, ABDUA:10739 and C14262, ABDUA:10742) the best-fitting model suggests a substantial proportion of ancestry that best fits *Iron Age Scandinavia*: 21±8%; *p*= 0.614 and 23±8%; *p*=0.08, respectively (**Fig. 1d**). While based only on two individuals, this result suggests contacts between Pictish-era Scotland and Migration-era Scandinavians predating the arrival of Vikings in 793 CE. Links to Scandinavia have also been suggested at a different Pictish site, Portmahomack, based on isotopic analysis^73^, and may explain the presence of ancestry related to Iron Age Britain in Viking Age northern Scandinavia^21,23,74^

Historical accounts have highlighted possible instances of matrilineality when describing royal succession amongst the Picts^75^. We note that two mitochondrial lineages (Jb1a1a and J1c3b) were detected at ∼17% frequency in Lochhead Quarry (with J1c3 also present at high frequency in another Pictish site, Lundin Links^71^), which could reflect matriliny-associated structure, but we caution that aDNA datasets from Pictish-associated contexts remain too small for a reliable population-level assessment of mitochondrial variation to exclude that this the result of genetic drift. Notably, none of the individuals in Lochhead Quarry were related to each other (up to 6th degree), but we identify a distant relative (up to 6th degree) of the Rosemarkie individual (441–641 cal. CE^76^) in the Isle of Iona, ∼200 km from Rosemarkie cave (C12388, undated) (**Extended Data Table 6**). This finding is in agreement with Morez et al.^71^ who suggested that Pictish burial practices may not be closely correlated with biological kin. Taken together with our new data from Lochhead Quarry this opens the possibility that life-time mobility may have been common.

We also inferred low impact of continental ancestries in sampled early medieval sites from present-day Wales, Somerset, Dorset and Cornwall (**Fig. 1d**; **Fig. 3; Supplementary Fig. 1**). Only a small number of individuals from these sites carried continental ancestries at variable proportions, including: C14205 (Sk 15) and C14207 (2C5.3), buried in a possible execution cemetery in the mound and ditch at Wor Barrow; three individuals from an undated mass grave in Portbury School C10680 (Box 4, Bag B2), C10682 (Box 2, Bag B) and C10683 (Box 1, Bag Cb); and one individual, C11655_prov (3081), from Yatton – all located near or outside the borders of archaeological evidence of early continental influence. The results from Portbury and Wor Barrow may be examples of conflict between different social and/or cultural groups, although the same sites also include individuals whose ancestry was entirely derived from Iron Age Britain. In present-day Wales, we detect disparate ancestries in Five Mile Lane (C12713, C11340, C11337, C12708, C12707, C12702) across different time periods, and in a 8th-11th century CE Christian cemetery at St. Patrick’s Chapel (C12684, C12686, C12690, C10655, C10660), in agreement with archaeological links between the Irish Sea region and continental Europe^77,78^. These findings highlight that while there were clearly factors inhibiting the flow of continental ancestry into western Britain, there were people with complete or partial ancestries derived from continental Europe in regions that were not predominantly Germanic-speaking.

Although *qpAdm* modelling points to apparent ancestry continuity in both northern and western regions throughout the 1st millennium CE, we caution that we do not formally test any ancestry sources from Ireland due to an absence of relevant published shotgun data. Contacts across the Irish Sea were a feature of the post-Roman period, as reflected in the linguistic landscape of early medieval western Scotland ^70^, but are unaccounted for in our current models. We also note in some cases the continental ancestries detected might be due to links to the Viking world (**Fig. 4**). The clear absence of major continental ancestries is also reflected in a general absence of IBD-segments shared with continental Europe.

### Changing mobility reflected in long-haplotype sharing patterns

We conducted large-scale IBD analysis of pairs of individuals, including also previously published individuals with only SNP capture data available^27^ (**Extended Data Table 6**). To account for the non-uniform nature of the published ancient record, we report IBD-sharing with continental individuals post-dating the Iron Age (instead of restricting the continental individuals to the early medieval period). By focusing on long IBD segments (>25 cM), we detected recently-shared ancestry between Britain and continental Europe, with striking geographical and temporal patterns. Individuals from Britain dating to the 5th-8th century CE shared more long IBD segments with individuals from continental North Sea coast, present-day north Germany and south Denmark, as well as with Longobard-associated individuals in present-day Hungary^79^ (**Fig. 4a**). These results are further supported by the aforementioned pair of distant relatives found in Cambridgeshire and Frisia.

By contrast, despite the *qpAdm* models pointing to substantial presence of ancestries related to Central Europe from the mid 8th century CE in Britain, the main IBD connections are instead with individuals from Viking Age Scandinavia, with an overall absence of IBD-sharing with currently available individuals from elsewhere in continental Europe (with the exception of a connection with an individual from the aforementioned Langobard context in Hungary) (**Fig. 4a**).

We also identified long IBD-segments shared between individuals in Britain and Ireland. Two female burials from Martyr’s Bay, Iona, Scotland, Lot 13 (C12397) and Lot 25 (C12395), both modelled with 100% Iron Age Britain ancestry, shared elevated IBD with a young male (VK543; EP55)^23^ found in a Viking grave in Eyrephort, County Galway, and with KIL041^6^, Burial 116 from a 6th-12th century CE burial ground at the Bishop’s Seat, Kilteasheen, County Roscommon, respectively^23^, as well as one individual from the Viking-associated Ridgeway Hill Mass Grave in Dorset (VK263)^23^, also displayed high IBD with individual VK543 (EP55). Although these results seem to point to a connection with Viking-related contexts, considering the low number of genomes available from Iron Age and medieval Ireland and from (north)west Britain, it is likely that our IBD analysis is currently underestimating connections across the Irish Sea.

IBD-sharing is highly stochastic and sampling-dependent, therefore caution is needed when interpreting these geographical patterns. Individuals sharing such long IBD segments most likely shared a common ancestor within the last 10-20 generations^80,81^ (with longer segments representing more recent shared ancestors), but that does not imply that they were directly, or ‘vertically’, related. Alternatively, their common ancestor could have lived in a different location to the identified individuals, possibly in a region unrepresented in our dataset. Similarly, the pair of distant relatives may have lived contemporaneously, or have been separated by up to six generations. IBD does not offer any indication of directionality of ancestry influx or extent of admixture. Nevertheless these IBD-sharing patterns reflect shared ancestry amongst individuals associated with groups who likely spoke Germanic languages^22^ and signal movements between continental Europe and Britain, as well as connections between Britain and Ireland (although in the later case more sampling will be needed to uncover any temporal and geographical patterns).

### Genetic impact of the Viking Age

Following the first recorded Viking raids at Portland, on the south coast of Britain, in c.789 CE and the monastery of Lindisfarne, in the northeast, in 793 CE^82^, the arrival of Scandinavian raiders and settlers had profound impacts on the social and political organisation of Britain (**Fig. 1a**), leading to the fracturing of the kingdoms of Mercia, Northumbria and East Anglia, and the increase of power and influence of the kingdom of Wessex (including the annexation of Cornwall, Dorset and Somerset). This culminated in the establishment of the Danelaw, and eventually the annexation of England to the North Sea Empire of the Danish Kings Svein and Cnut for approximately 30 years in the 11th century CE^64^. In northern and western Britain, many of the islands and coastal zones, including the Isle of Man, would fall to Scandinavian control, with trade and mobility now dominated by the emerging Earldom of Orkney and the early Hiberno-Scandinavian trade settlements across the Irish Sea^11,83^.

While ancestry was diverse in Viking Age Scandinavia^21,23,74^, the patterns of ancestry seen in the late 8th-11th centuries CE in Scandinavia and Britain differ substantially (**Fig. 4c**). In Scandinavia (Denmark, Norway and Sweden), 31-65% of individuals dating to 750-1050 CE retain ancestry best modelled by Iron Age Scandinavia, compared to only 4% in England (**Fig. 4d)**. To further examine this, we used a Chromopainter-like approach based on shared recent coalescences between individuals, summarised using non-negative least squares (NNLS) (**Methods**). We then assessed the resulting dimensions for features that separate individuals in Britain with clear contextual links to the Viking world, from others in the same period. We identify two distinct clusters dominated by previously-published individuals from Viking-associated contexts in Oxford and Dorset^23^, as well as newly sequenced individuals mostly from contexts showing Scandinavian influences or associated with Viking contexts (**Fig. 4b**): two individuals from a cemetery in Mayback, Orkney, including a female individual from a boat burial radiocarbon dated to 772-885 cal. CE (C12546, Sk 012); C15255 (Sk 4), from the Red House Park Sewer site, Adwick−le−Street in South Yorkshire, who was accompanied by two classic Scandinavian oval brooches^84^; C12544_prov (Sk003) from the graveyard of a 10th century CE chapel at Newark Bay; two individuals from a 9th-10th century cemetery adjacent to the 872/873 CE Viking Great Army winter camp in Torksey, Lincolnshire^85^; C12607 (Sk 14), from a 8th-13th Century CE cemetery without any archaeological evidence of Viking contacts in Barnstaple Castle, Devon; and C10138, a burial radiocarbon dated to 990-1036 cal. CE from St. Ninian’s Chapel, in Bute, western Scotland.

The association between these distinctive clusters and sites or individuals showing archaeological connections to Scandinavia suggest that we are detecting subsets of ancestries associated with Viking Age raiders or settlers. However, given the high genetic diversity observed amongst these individuals (**Fig. 1d**), mirroring that observed in Scandinavia during this period (including similar influx of Central Europe-related ancestries into southern Scandinavia; **Fig. 4c**), this could represent an underestimation and/or a failure to identify individuals with admixed ancestries. Overall, only 31 of 226 individuals in this period are distinct in PC3 but this is likely an overestimation due to our and previous^23^ sampling targeting Viking-associated contexts. Together with the Twigstats ancestry models, this suggests that the population-level impact of the Viking Age was more modest than that of the Migration Period in Britain.

### Ancestry-specific single-locus selection

Previous aDNA studies focussing on selection in West Eurasia over the past 10,000 years often lacked the resolution to link selection to particular periods or regions^7,29,31^, using statistical models to account for broad ancestry differences, which may have missed subtle ancestry confounders that could obscure allelic changes resulting from adaptive evolution. In the case of Britain, previous studies have detected selection on *LCT* and *DHCR7* associated with diet and Vitamin D metabolism, *SLC45A2* and *HERC2* associated with pigmentation, and the HLA region associated with immunity^28,30^, but without accounting for the diversity of ancestries in the medieval period reported here.

To detect signatures of natural selection in a conservative time series with maximal ancestry continuity, we used our dense time series of 427 individuals showing ancestry continuity to test for a significant association between derived genotype frequencies through time using linear regression^86^, on a compiled set of imputed whole-genomes data on ∼5.5 million variants, using genomic control to adjust *p*-values for genome-wide temporal drift (**Methods**). Specifically, we restricted the test to individuals dating to the Bronze and Iron Ages (after removing outliers; **Extended Data Fig. 5**) and later individuals (1-1150 CE) that were consistent with deriving 100% of ancestry from Britain Iron Age based on our fine-scale Twigstats-*qpAdm* ancestry modelling (*p*>0.01) (**Fig. 5a**). This approach removes all detectable confounding effects deriving from transient mobility in the Roman period and large-scale ancestry transformations in the early medieval period. While we do not explicitly correct for ancestry transformation in the Middle to Late Bronze Age^10^ (**Extended Data Fig. 4a)**, we corroborated the results using a mixed linear model with the Genetic Relationship Matrix (GRM) to control for any remaining effects of population structure (**Methods; Extended Data Fig. 6a**).

**Fig. 5:**
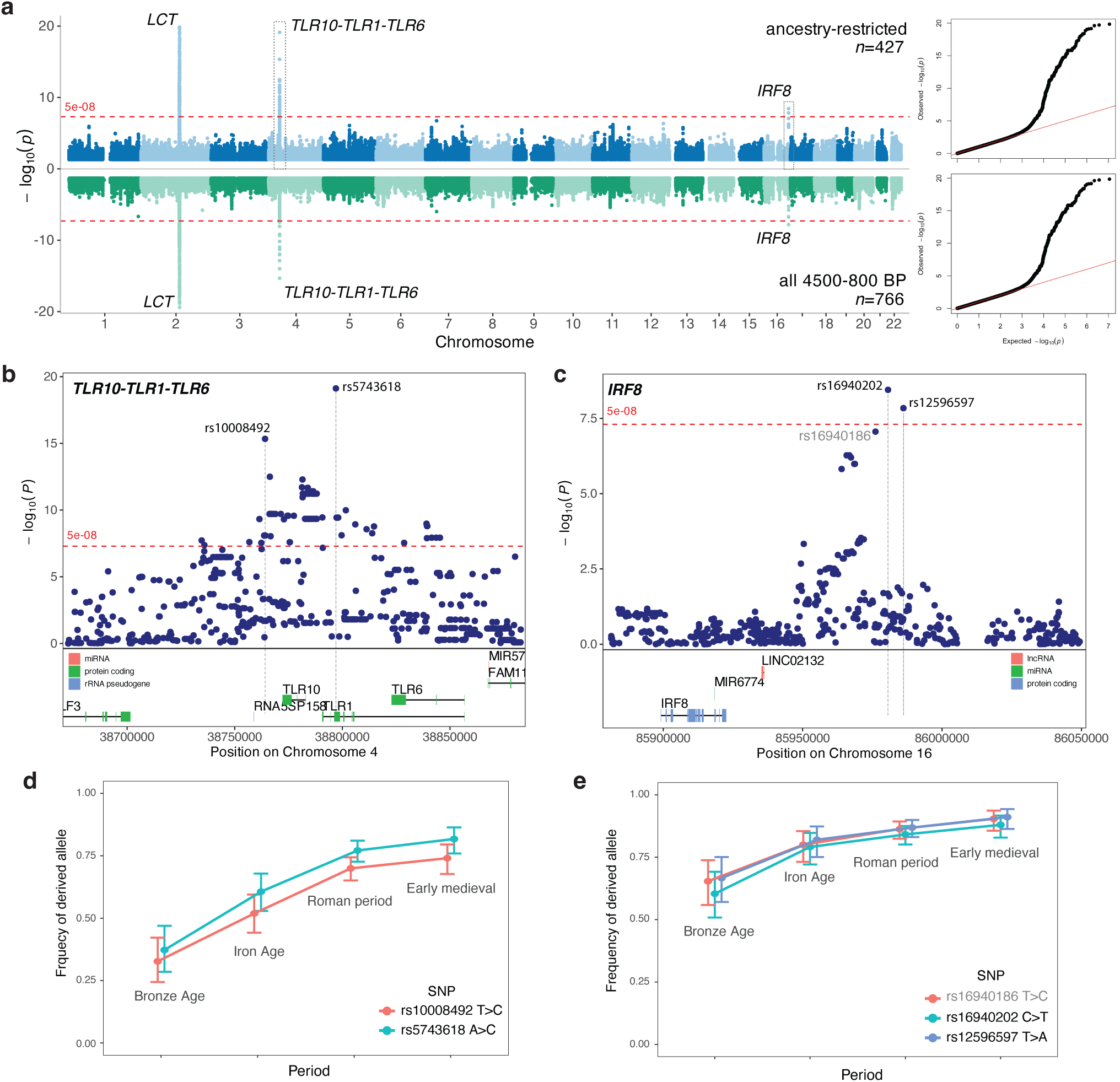
Ancestry-restricted selection scan. **a)** Manhattan plots of *p-*values and QQ plots from linear regression with genomic control on two datasets: ancestry-restricted dataset (**Methods**) in blue (*n*=427) and dataset including all individuals dating to 4500-800 BP (after excluding obvious outliers, *n*=766; Extended Data Fig. 6) in green. Closest protein-coding genes are labelled next to significant hits. **b)** Zoom in the TLR10-TLR1-TLR6 region on chromosome 4. **c)** Zoom in the region around *IRF8* on chromosome 16. **d)** Allele frequencies of the top two SNPs on chromosome 4 (rs5743618 and rs10008492) in Britain in different archaeological periods. **e)** Allele frequencies of the top three SNPs on chromosome 16 (rs16940202, rs12596597 and rs16940186) in Britain in different archaeological periods.

We detect the previously described selection signal on *LCT*^28^. While the major allele frequency increase was prehistoric^10,28^, our large-scale sampling of the Roman period and ancestry-stratification of individuals suggests that the frequency may have continued to increase in the first millennium CE (**Extended Data Fig. 6b**).

We find a strong signal of allele frequency change in the *TLR10-TLR1-TLR6* region with rs5743618 and rs10008492 as the two top SNPs (*p*=7.6e-20 and 4.6e-16, respectively, *n*=427) (**Fig. 5b**). SNPs in this region were identified in an early pan-European selection scan between the Bronze Age and present-day^29^, and rs10008492 was just below standard significance thresholds in a previous scan of Britain that did not account for ancestry change (*p*=10^-7^, *n*=504)^30^. The derived allele frequency increased in the Bronze Age (**Fig. 5d**), but also notably between the Iron Age and the Roman period, from ∼50% to ∼70%. rs5743618 is located in a protein-coding region of *TLR1*, whereas rs10008492 is near a regulatory region active in most primary immune cells with the notable exception of T-cells (**Extended Data Fig. 6d**), and this haplotype has been specifically associated with increased risk of hayfever, allergic rhinitis or eczema^87^ and with higher C-Reactive Protein (CRP) levels, an acute phase inflammatory marker^88^. The derived allele frequency of rs5743618 also increased between the Iron Age and the Roman period from ∼60% to ∼77%. Both SNPs are in linkage disequilibrium (LD) (*r*^2^=0.57 in 1000 Genomes Project GBR) and are rare outside modern populations with European-related ancestry (**Extended Data Fig. 6c**).

*TLR1* and *TLR10* encode pattern recognition receptors that are involved in the innate immune response to microbial pathogens, but with opposite effects: *TLR1* is inflammatory and *TLR10* is anti-inflammatory. The *TLR1* SNP is a derived missense coding variant (rs5743618 - I602S) showing signatures of positive selection in modern European populations^89^, and inhibits the surface trafficking of the TLR1–TLR2 heterodimer, reducing NF-κB activation upon ligand recognition^90^. The dual signal of selection of the I602 missense variant in *TLR1* suggests that availability of TLR1 is also reduced. In contrast, the enhancer-linked rs10008492 variant is significantly associated with increased expression of TLR1 (*p*_FDR_=4.39e-7) and decreased expression of TLR10 (*p*_FDR_=0.02) in stimulated macrophages; effects that in combination would enhance inflammatory responses (notably no significant effect was observed in unstimulated macrophages; **Extended Data Fig. 6e**). One possible unifying explanation is that the variants in concert decrease TLR10 and TLR1 binding to TLR2. This would free up TLR2 to bind to TLR6 instead. Whereas the TLR1-TLR2 heterodimer recognises triacylated lipopeptides common in gram-negative bacteria, the TLR6-TLR2 heterodimer–for which we find putative selection for increased availability–recognises diacylated lipopeptides common in gram-positive bacteria such as *Streptococcus* and *Staphylococcus*. Further evolutionary and functional research will be able to test this hypothesis.

In addition, we found two marginally significant SNPs on chromosome 16, rs16940202 and rs12596597 (*p*=6.441e-10 and 2.997e-09, respectively; plus a third one just under the threshold of significance, rs16940186; *p*=2.097e-08), near *IRF8* (**Fig. 5c,e**), which encodes the interferon regulatory factor (IRF)8. IRF8 is a transcription factor involved in the development of myeloid cells (including manocytes and macrophages), expressed mainly in the spleen and lymph nodes, but also in the intestine and heart^91^. SNPs within or in the vicinity of *IRF8* impact blood cell count for monocytes, neutrophils and eosinophils, with a postulated impact on susceptibility to infections^92,93^. IRF8 expression in stomach and intestinal epithelium cells has been linked to resistance to gastrointestinal infections, whereas IRF8 deficient gastric cells fail to induce inflammatory responses during infection leading to higher bacterial loads and increased tissue damage^94^. On the other hand, genome-wide association studies (GWAS) have shown an association between SNPs in this region and chronic inflammatory diseases, with rs16940202 and rs16940186 both linked to ulcerative colitis and inflammatory bowel disease^95–97^.

Together, these results emphasise the key role of infection in this period in Britain. For the *TLR* loci notable change occurs in the ∼200 BCE to ∼400 CE period, coinciding with increasing population density, emerging urbanisation and greater long-range mobility facilitated by the Roman Empire. Combined, these factors could have facilitated the spread of infectious diseases that modulated the inferred selection processes – including the Antonine plague epidemic. Future data will provide opportunities to test these timings and infer driving selective pressures in greater detail.

## Conclusions

Shifting geopolitical processes and their implications for aspects of social kinship during the 1^st^ millennium CE had contrasting impacts on the genetics of populations in Britain through time. Rome’s annexation of a large part of Britain (43-c.410 CE) was associated with increased incidence of individuals with novel ancestries in urban, elite and military centres, but the broader population showed substantial genetic continuity from the Iron Age. Nevertheless, our results suggest an increase in insular mobility during this era, with Roman administration and cultural norms possibly leading to changes in the formerly regionally-focussed patterns of inheritance and relations.

In contrast, we see a large-scale transformation of genetic ancestries in southern and eastern Britain from at least the 6th century CE, reflecting movements of people from across the North Sea^59^. These results parallel population dynamics inferred across parts of continental Europe^21,22,65^, consistent with a scenario of increased mobility into Britain and admixture in the era of western Roman imperial dissolution. It is possible that some of these population dynamics were either related to, or triggered by, the fall of the western Empire. The heterogeneous temporal and geographical patterns of continental ancestries detected here in early medieval Britain could represent different waves of migration or continuous migration (the latter supported by recent isotopic results^59^), with the cremation rituals of the 5th century CE rendering part of this population shift invisible in the aDNA record. Future isotope work on cremated remains has the potential to shed light on this question^98^. In parts of northern and western Britain we infer a much lower impact of these incoming ancestries, but quantification of the likely impact of connections to Ireland is still hampered by the lack of comparative data.

The substantial continuity in ancestry from the Iron Age into the Roman period in Britain, and the retention of this ancestry in northern and western regions through the early medieval period, allowed us to build an ancestry-homogeneous dataset and directly pinpoint strong evidence of natural selection in immunity within a couple of millennia, without resorting to extrapolations into the past or from admixed individuals. This study showcases the potential of large-scale ancient shotgun genome datasets to achieve a high-resolution view of ancestry and relatedness at different scales, providing more nuanced insights into population dynamics over time, and their role in untangling fine-scale evolutionary history.

## Data availability statement

New genomic data analysed in this project will be made available at the European Nucleotide Archive.

## Acknowledgements

This work was supported by a Wellcome Trust Investigator Award to P.S. (217223/Z/19/Z), and P.S. was also supported by a UKRI Horizon guarantee/ERC Consolidator award (UKRI338), the European Molecular Biology Organisation, the Vallee Foundation, the European Research Council (grant no. 852558), and Francis Crick Institute core funding (FC001595) from Cancer Research UK, the UK Medical Research Council, and the Wellcome Trust. L.K. received funding from Badenoch and Strathspey Heritage (Badenoch Storylands, National Lottery Heritage Fund Great Place Scheme grant) and Findlay Harris Dick Prize for Pictish Research. S.L. and K.L.S. acknowledge a Leverhulme Trust Early Career fellowship award (ECF-2021-467). S.L. was also supported by Cambridge Trust and Newnham College Studentship No. 10386281. L.K and K.Br. were supported by the Leverhulme Trust (Comparative Kingship project, RL-2016-069) and the University of Aberdeen. K.Br. was further supported by Leverhulme Trust (PLP-2019-284) and ERC-selected/UKRI-funded award (EP/Y023641/1). O.R.A. acknowledges support from Homes England. C.Booth acknowledges Persimmon Homes Severn Valley. M.G.A. acknowledges support from National Grid and Balfour Beatty. E.B. and M.G. acknowledge Skanska on behalf of the Defence Infrastructure Organisation (DIO). M.H. acknowledges the Society of Antiquaries of Scotland. R.J.S. acknowledges the Oxford Radiocarbon Accelerator Unit. A.J.E.P. and L.H.E. acknowledge the Roman Research Trust (RRT). G. N. was supported by Badenoch and Strathspey Heritage (Badenoch Storylands, National Lottery Heritage Fund Great Place Scheme grant) and the Leverhulme Trust (Comparative Kingship project, RL-2016-069). I.A. received funding from the European Research Council (ERC), under the European Union’s Horizon 2020 research and innovation programme, grant agreement no. 834087 (COMMIOS). M.G.T. is supported by ERC Horizon 2020 research and innovation programme grant agreements: no. 951385 (COREX) awarded to M.G.T., no. 865515 (SUSTAIN) awarded to Maria Ivanova-Bieg, no. 324202 (NeoMilk) awarded to Richard Evershed, no. 788616 (YMPACT) awarded to Volker Heyd, by Wellcome Senior Research Fellowship Grant 100719/Z/12/Z awarded to M.G.T, and by UK Natural Environment Research Council ‘Pushing the frontiers of environmental research’ grant NE/X01469X/1 awarded to M.G.T. J.C.L is a Lister Institute Prize Fellow and was supported by Francis Crick Institute core funding (CC2219). L.S. acknowledges JSPS KAKENHI grant 24K23946. We thank Andrew Armstrong, James Arnold, Mark Atkinson, Megan Beale, Oliver Blackmore, Matthew Bland, Katherine Buckland, Ciara Butler, Andrew Chamberlain, Barry Chandler, Sarah Chubb, Coralie Clover, Glynn Davis, Jodie Deacon, Natasha Dodwell, Paula Gentil, Adrian Green, Martin Green, Philip Hadland, Emma Harper, Mike Henderson, Neil Holbrook, Petra Ivanova, Elizabeth Johansson-Hartley, Bernard Jones, David Kaye, Tom Lord, Andy Maxted, Alison Mills, Graham Mullan, Ken Murphy, Benjamin Neil, Richard Osgood, Jenny Oxley, Adam Parker, Iwan Parry, Joe Perry, Louise Rayner, Mark Redknap, Andrew Reynolds, Andrew Richardson, Peter Robinson, James Sainsbury, Yulia Shevlyakova, Andrew Simmonds, Siobhán Sinnott, Agata Socha-Paszkiewicz, Allan Steward, Gabor Thomas, Brett Thorne, Ross Turle, Clive Waddington, David Walker, Ros Westwood, Linda Wilson, John Wood, Andrew Woods, Jenifer Woolcock and Tom Woolhouse, as well as Amgueddfa Cymru, AOC Archaeology, Archaeological Research Services Ltd, Avon Archaeology, Buxton Museum and Art Gallery, Cambridge Archaeological Unit, Department of Archaeology, University of Cambridge, Canterbury Archaeological Trust, Canterbury Christchurch University, Colchester Archaeological Trust, Cotswold Archaeology, Cura Terrae, Discover Bucks Museum, Dover Archaeological Group, Dover Museum, Down Farm Museum, Dorset, Eastbourne Borough Council, Folkestone Museum, Foundations Archaeology, Hastings Museum and Art Gallery, Heritage Doncaster, Historic England, Hull and East Riding Museum (Hull Museums), KDK Archaeology, Levens Local History Group, Mill Green Museum, Museum of Barnstaple and North Devon, Museum of Gloucester, Museum of London Archaeology (MOLA), Museum of the Great Western Railway, Museum & Art Swindon and Lydiard, National Museums Scotland, Network Archaeology, Newport Museum and Art Gallery, Northamptonshire Archaeological Resource Centre, Oxford Archaeology Ltd, Pre-Construct Archaeology Ltd, Red River Archaeology, Royal Institution of Cornwall, Rubicon Archaeology Group Ltd, Saffron Walden Museum (Uttlesford District Council & Saffron Walden Museum Society), SLR Consulting Ltd, South West Heritage Trust, Thames Valley Archaeological Services Ltd, The Beacon Museum, The Hampshire Cultural Trust, The Harris Museum, The London Museum, The Potteries Museum and Art Gallery, The Salisbury Museum, The University of Bristol Spelaeological Society, Torquay Museum, University of Aberdeen Collections, University of Aberdeen School of Geosciences, University of Exeter, University of Leicester Archaeological Services, University of Sheffield, Wardell Armstrong, Wells Museum and Art Gallery, Wessex Archaeology, Wiltshire Museum, Devizes, Worthing Museum, York Archaeology and York Museums Trust for permission to sample individuals and access to collections. We thank Ken Lymer (Cotswold Archaeology) for providing the site plan shown in Extended Data Fig. 3. We thank Anastasia Brativnyk, Iain Mathieson, Lluis Quintana-Murci, Caetano Reis e Sousa, Oded Rimon, Maxime Rotival, Mattias Sherman and Eirini Skourtanioti for helpful comments and discussions on the genetic analysis, and Hella Eckardt and Martin Carver for valuable comments on an earlier version of this manuscript. We thank Ricardo Rodríguez-Varela and Anders Götherström for data sharing, Mattias Meyer for help with high-throughput robotic workflows, and Ferdinando Verdirame, the Genomics STP and Scientific Computing at the Francis Crick institute for technical support. For the purpose of open access, the authors have applied a CC BY public copyright licence to any Author Accepted Manuscript version arising from this submission.

## Author contributions

M.S., T.B. and P.Sk. designed the study. M.S., T.B., J.M., M.M., S.J., F.T., P.Sw., J.P., S.L., L.J.K., H.F., B.V.T., E.B-K., A.J.E.P., I.B. and S.B. performed sampling for DNA extraction and/or radiocarbon dating. M.S., K.A., M.K., M.W., S.J., F.T., P.Sw., I.G., M.R.C., I.B. and S.B. conducted lab work for DNA retrieval and/or sequencing. M.S., K.A., C.Ba., Y.M., A.G., M.R.C. and L.S. processed and/or curated sequencing data. M.S., K.A., J.C.L., L.S. and P.Sk. conducted analyses. T.B., J.M., M.M., S.L., L.J.K., E.B-K., O.R.A., M.G.A., E.B., C.Booth, C.Boston, A.B., L.B., C.C., E.C-A., N.G.W.C., A.J.D., G.D., J.d.P., L.H.E., C.F., M.G., D.G., K.A.H., A.H., M.H., A.N.R.H.K, M.G.K., S.M., K.P., A.J.E.P., R.R., J.S., C.A., S.S., M.C.S., A.C.T, J.E.T, D.W., G.W., H.W., A.W., A.Y., S.J.C., A.S., M.A.G., L.L., M.R.H., R.J.S., K.Br., K.Bu., G.N., I.A., J.B., I.B., M.G.T., L.G-F. and A.M. curated archaeological samples and/or provided archaeological contextualistion. T.B., S.L., L.J.K., E.C-A., D.G., M.R.H., R.J.S., K.Br., G.N., I.A., L.G-F., A.M. and P. H. provided historical and archaeological interpretation. L.G-F. supervised L.K., P.Sk. supervised the study. M.S and P.Sk. wrote the original draft. T.B., M.M., S.J., S.L., L.J.K., E.C-A., D.G., M.A.G., M.R.H., R.J.S., K.Br., G.N., I.A., J.B., I.B., S.B., M.G.T., L.G-F., A.M., P. H., J.C.L. and L.S. reviewed and edited the final draft.

## Supplementary Information

This PDF file contains Supplementary Figures 1 to 9:

**Supplementary Fig. 1:** Twigstats-boosted *qpAdm* models for individuals from Britain dating to the 1st millennium CE (as in Fig. 1d), with site labels for all targets and sample identifiers for individuals highlighted in the main text. Both newly sequenced and previously published (site labels in grey) individuals are included and are plotted according to their average date (**Extended Data Table 1**); note that the individuals included in the *Early Medieval Britain I* source (all dated to the 6th century CE) are also plotted. Only individuals with accepted *qpAdm* models are shown (*p*-value>0.01; **Methods**). Full *qpAdm* results in **Extended Data Table 8**.

**Supplementary Fig. 2:** Mitochondrial haplogroup frequencies in Iron Age sites with n≥5 (excluding close relatives).

**Supplementary Fig. 3:** Mitochondrial haplogroup frequencies in Roman-period sites with n≥5 (excluding close relatives).

**Supplementary Fig. 4:** IBD-sharing between individuals in Britain and outside of Britain in the **a)** Bronze Age, **b)** Iron Age and **c)** Roman period. Sites with one or more individuals sharing at least one IBD segment >25 cM with another individual in a different site outside of Britain are represented by circles, and both sites are connected and coloured according to total IBD sharing (>8 cM). Remaining sites included in the analyses are represented by a single red ‘x’. Dataset includes shotgun and capture genomes (**Extended Data Table 6**). Note that these maps do not show IBD-sharing within Britain or between pairs that do not include one individual from Britain.

**Supplementary Fig. 5:** Unsupervised ADMIXTURE clustering, from *K*=2 to *K*=15.

**Supplementary Fig. 6:** Twigstats-boosted MDS and qpADM modelling for early medieval and later targets in different periods: **a)** 410-600 CE, **b)** 600-750 CE, **c)** 750-1066 CE **d)** post-1950 CE. MDS based on Twigstats-boosted pairwise outgroup-*f*3s (*t*=1000 cutoff), stratified by time. Individuals from Britain (both newly sequenced and published) are used as targets and represented by their *qpAdm* modelling proportions (as in Fig. 2; showing only plausible and non-rejected models, i.e. *p*-value>0.01 and ancestry proportion±SE between 0 and 1). Number of targets (*n*) for each time period indicated on each MDS. Polygons represent source populations included in the *qpAdm* models (Sarmatian not included in the MDS) and previously published continental populations dating to the Roman and early medieval periods. Maps show the proportion of ancestry derived from each tested source for each individual. Model selection detailed in **Methods**. All *qpAdm* results in **Extended Data Table 8**.

**Supplementary Fig. 7:** Comparison of autosomal (verifyBamID) and X-chromosome (ANGSD) contamination estimated for libraries with a XY karyotype (**Methods**). Individuals with average autosomal coverage (MQ>30) under 0.1x coloured in dark red.

**Supplementary Fig. 8:** Imputation accuracy (*r*^2^) for different downsampled coverages (5x, 2x, 1x, 0.5x, 0.3x and 0.1x) at different maf, comparing imputation with and without applying baSSdrop damage correction for 12 high-coverage samples generated using ssDNA library preparation (**Methods**). Initial coverages (hcov) varied 7.5–10.3x after remapping to GRCh38 and retaining only reads with MQ>30.

**Supplementary Fig. 9: a)** Sum of IBD1 >20 cM segments and total length of genome in IBD1 (>20 cM). **b)** Correlation in total length of IBD1 >12 cM identified with ancIBD and KIN (r2=0.96). Pairs are coloured according to relatedness as estimated with KIN (for 1st-3rd degree); 4th-6th degree relatives annotated for pairs sharing at least two independent IBD-segments >20 cM, following Ringbauer et al. Complete relatedness results shown in **Extended Data Tables 6 and 7.**

–--

This Excel file contains the following 13 Extended Data Tables:

**Extended Data Table 1:** Archaeological information and sequencing metrics for 1039 individuals with whole shotgun genomes generated in this study.

**Extended Data Table 2:** Mitochondrial diversity indices in Iron Age and Roman-period sites.

**Extended Data Table 3:** Compiled information on matrilineal burial practices and consanguinity for all Iron Age sites.

**Extended Data Table 4:** Statistical tests supporting association between finding a consanguineous individual and a matrilineal burial signal in the site. **a)** One-sided Fisher’s test showing a significant association between finding a consanguineous individual and matrilineal burial signal in an Iron Age site. Classification of matrilineal burial signal as reported in Extended Data Table 2. **b)** Kruskal-Wallis rank sum test showing a significant association between ROH in 4-8 cM bins (**b1** and **b2**) and total ROH > 4cM (**b3**) and matrilineal burial signal in Iron Age sites.

**Extended Data Table 5:** Number and length of runs of homozygosity (ROH) for individuals sequenced in this study carrying at least one ROH>4cM.

**Extended Data Table 6:** Identity-by-Descent (IBD) shared between all pairs included in the dataset, excluding pairs of close relatives (shown in Extended Data Table 6).

**Extended Data Table 7:** Identification of close relatives (up to third degree) in the newly-sequenced dataset, using both ancIBD and KIN. For pairs with a KIN Log Likelihood Ratio (LLR) <1 we also report the second most-likely inferred relatedness (if third degree or below) or add’?’ if second inferred relatedness is above third degree.

**Extended Data Table 8:** Twigstats-boosted (*t*=1000) *qpAdm* modelling for individuals from Britain (both newly sequenced and previously published) and relevant individuals from continental Europe (previously published) dating to the 1st millennium CE. All models with *p*-value>0.01 are included (even if with implausible ancestry proportions plausible). Model adapted from Speidel et al. 2025 (Methods).

**Extended Data Table 9:** Twigstats-boosted (*t*=1000) *qpAdm* modelling for Roman-period individuals from Britain (both newly sequenced and previously published), including an additional Iron Age Britain (post-200 BCE) source. All models with *p*-value>0.01 are included (even if with implausible ancestry proportions plausible). Model adapted from Speidel et al. 2025 (Methods).

**Extended Data Table 10:** QC metrics for new radiocarbon dates generated in this study.

**Extended Data Table 11:** Distal *qpAdm* model with fixed sources – Western Hunter-Gatherers (WHG), Early European Farmers (EEF) and Steppe, based on Patterson et al. 2022 – for newly sequenced individuals (pseudo-haploid genotyped for ‘1KGP transversion sites’; Methods). Rejected models with *p*<0.01 highlighted in red.

**Extended Data Table 12:** Twigstats-boosted (*t*=5000) *qpAdm* model with fixed distal sources – Western Hunter-Gatherers (WHG), Early European Farmers (EEF) and Steppe, based on Patterson et al. 2022 – for a subset of 878 newly sequenced individuals that were included in the Relate run. Rejected models with *p*<0.01 highlighted in red.

**Extended Data Table 13:** Chromopainting datasets, with list of individuals included as targets and palette for each period: **a)** Iron Age, **b)** Roman period, **c)** early medieval.

## Methods

### Archaeological sampling for aDNA and ethics framework

Sampling was performed in clean room facilities at the Francis Crick Institute and at the Ancient DNA laboratory at the University of Aberdeen, or ‘on site’ whenever necessary. We followed the guidelines issued by the Department for Culture, Media and Sport (DCMS) and the Advisory Panel on the Archaeology of Burials in England (APABE) for minimally-destructive sampling (https://apabe.org.uk/resources/documents-apabe).

For crania and loose temporal bones we either collected single auditory ossicles (retrieved from the auditory canal)^101^ or collected multiple subsamples of bone powder from the cochlear portion of the petrous bone^102^ by drilling a single ∼1-4 mm wide hole from the jugular fossa using an Emax EVOlution (EV410) micro-motor system with disposable carbide round burs. For teeth, we collected a cementum-enriched subsample, by scraping the surface of single roots, and/or drilled a single ∼1-3 mm wide hole into the pulp chamber to collect subsamples of dentine powder. In addition, we also shotgun-sequenced genomes of 12 individuals sampled elsewhere and previously subjected to the 1240k enrichment protocol. The final dataset comprises 1039 individuals, sequenced from 594 single ossicles, 351 powders from the cochlear proportion of the petrous bone, 85 teeth, and, for a small proportion of individuals (*n*=9), combined multiple elements.

Our sampling reflects a wide variety of archaeological contexts. Although we tried to maximise geographical, as well as temporal, representation, regions of acidic soils with poorer bone preservation^103^ and/or lower-density human occupation with fewer opportunities for archaeological surveys (Devon and Cornwall, the interior of Wales, and the northwest of England and Scotland, in particular the Highlands), as well as periods of widespread cremation or other funerary rites that leave little archaeological trace^53,60^ (*e.g.* excarnation, riverine depositions) remain undersampled (**Fig. 1a**). For the Roman period, we note that the sampling covering the 50-200 CE period is significantly lower than that for the 200-410 CE period (*n*=55 vs *n*=257) – possibly reflecting the scarcity of inhumations as a a funerary practice in the early Roman period^53^ – and only ∼20% of these individuals come from three urban or military contexts more likely to harbour incoming ancestries. We also caution that there is a degree of uncertainty for late Roman inhumations that are often dated by association to artefacts and could in some cases be later, as demonstrated before^104^.

### Radiocarbon dating

We report 31 new radiocarbon dates (**Extended Data Table 10**). We sampled skeletal material from 21 individuals at the Francis Crick Institute, using an Emax EVOlution (EV410) micro-motor system with attached diamond disc rotary tool where relevant. Sample sizes of >500 mg were taken where possible, with a minimum threshold of 100 mg. For 14 samples we applied a bespoke collagen extraction, using the pretreatment method with ultrafiltration outlined in Fewlass et al. 2019^105^. The >30 kDa extracted collagen fractions were sent to either the radiocarbon facility at the University of Bristol (BRAMS) or the University of Oxford (ORAU) for graphitisation and AMS dating. Seven samples were sent directly to the radiocarbon facility at ORAU for collagen extraction and AMS measurements. We prepared two samples (ArchaeoFINS) in the Andy Barlow Bone Chemistry Laboratory, School of History, Classics and Archaeology, University of Edinburgh. Bone collagen and dentine were processed following a modified Longin method^106,107^, and sent to the Scottish Universities Environmental Research Centre (SUERC) to undergo EA-IRMS and AMS measurements.

We recalibrated (95.4% confidence interval) radiocarbon dates with OxCal v.4.4.4 using the IntCal20 calibration curve^108,109^ (**Extended Data Table 1**). For samples where carbon and nitrogen stable isotopes indicated the need to correct for a marine reservoir effect, this was done in OxCal v.4.4.4 using a mixed curve model^87,88^ with a marine endpoint of −12.5‰ and a terrestrial endpoint of −21.0‰ for δ13C and an ΔR value of-124±44^110^.

### DNA extraction and sequencing

DNA extraction and pre-amplification steps were performed in specialised clean rooms at the Francis Crick Institute, physically separated from lab areas handling modern samples or post-PCR products. We extracted DNA from single ossicles, or from ∼1-164 mg of bone/tooth powder subsamples^111^, and prepared double-indexed single-stranded (ss) DNA libraries^112,113^ without performing any UDG-treatment, using automated liquid-handling systems (Agilent Bravo Workstations), with the exception of 12 double-stranded (ds) DNA libraries that were previously prepared at the Natural History Museum for which capture data has been published before^6,10,114^ and we shotgun-sequenced here (**Extended Data Table 1**). Negative extraction and library controls, as well as positive library controls, were processed alongside samples.

To assess DNA preservation and authenticity, libraries (including negative controls) were initially screened using Illumina sequencing platforms (NextSeq 550, HiSeq 4000, NovaSeq 6000), and later selected for further sequencing based on the proportion of endogenous human DNA, evidence of 5’-and 3’-end C>T substitutions and library complexity. Selected libraries were further sequenced on Illumina NovaSeq 6000 or NovaSeq X platforms in different proportions based on screening metrics, in an attempt to maximise autosomal coverage for each sample. A subset of these libraries were subjected to a gel-excision protocol^112^ to retain only DNA sequences between 35–150 bp prior to sequencing (**Extended Data Table 1**), but this protocol was later discontinued after an initial assessment showed no substantial improvement on the final predicted coverages relative to sequencing effort.

### Initial sequencing data processing

Sequencing data was processed using nf-core/eager^115^ v.2.3.3^116^. We merged within-flow cell sequencing replicates, provided *Colour_Chemistry* arguments according to the Illumina platform, *SeqType* as ‘PE’, *Strandedness* as ‘single’ or ‘double’ and *UDG_Treatment* as ‘none’. For data generated on two-colour chemistry machines, the *--complexity_fiilter_poly_g* argument was also provided. Forward and reverse adapters were provided with *--clip_forward_adapter* ‘AGATCGGAAGAGCACACGTCTGAACTCCAGTCAC’ and *--clip_reverse_adapter* ‘GGAAGAGCGTCGTGTAGGGAAAGAGTGT’ (for ss libraries). A minimum read length of 35 bp was applied with *--clip_readlength* and the *--preserve5p* option specified to avoid trimming the 5’ end of reads. Only merged read pairs were considered for analysis using the *--mergedonly* argument. We aligned merged reads with bwa^117^ v.0.7.17-r1188 *aln* and *samse*, using the hs37d5 reference genome with default nf-core/eager arguments (-n 0.01-l 1024-k 2), and removed potential PCR duplicates with DeDup^118^ v.0.12.8.

For samples with more than one library we merged BAM files (samtools^119^ v.1.3.1 *merge*). When the same library was sequenced more than once, we subsequently removed duplicates (DeDup *-m*).

### Karyotypic sex and chromosome 21 karyotype

We estimated karyotypic sex (**Extended Data Table 1**) using a script that accounts for chromosomal aneuploidies^100^ by comparing the ratios of sequencing reads mapped to each sex chromosome relative to the autosomal baseline. We used the option *--chr21* to identify potential cases of trisomy 21.

### Data authenticity and contamination estimates

We estimated contamination in the X chromosome of libraries assigned as XY using ANGSD^120^ v.0.933, and assessed mtDNA contamination for all libraries using schmutzi^121^ v.1.5.5.5 (*contDeam.pl --library single* or *--library double*, depending on library protocol) (**Extended Data Table 1**). In addition, we ran VerifyBamID^122^ v2.0.1 for all libraries (with parameters *--WithinAncestry*, *--min-BQ 30* and *--min-MQ 30*) to estimate autosomal contamination by comparison with a database of genotypes from the 1000 Genomes Project (1KGP) phase 3 restricted to ∼206,500 transversion sites overlapping with the ‘1240k capture panel’. We further checked contamination by inferring karyotypic ratios^100^.

We considered libraries as contaminated when they met at least one of the following criteria (**Supplementary Fig. 7**): autosomal contamination >5% (estimated either with ANGSD for XY individuals, or VerifyBamID), mtDNA contamination >10%, or abnormal karyotypic ratios consistent with contamination. We flagged as contaminated one additional library displaying suspiciously low (<1%) C-to-T misincorporation at 5’-end of sequencing reads. We annotated as ‘provisional’ XX libraries with autosomal contamination estimates 3-5% (for which we do not have angsd estimates) or any library with schmutzi estimates 5-10%, as long as all other contamination estimates were below the thresholds mentioned above. We included all libraries in relatedness analyses, but we interpret the results with caution, and excluded contaminated libraries from all further analyses. Libraries labelled as ‘provisional’ were included in all analyses (providing they met the coverage thresholds required for specific analyses).

### Remapping to GRCh38, ss-aware damage correction and imputation

We remapped our newly sequenced genomes, as well as relevant published shotgun and ‘1240k’ data, against GRCh38 reference using bwa *aln* (using *-b* flag to specify BAM format as input, *-l 1024-n 0.01-k 2*) and *samse*.

To avoid potential imputation biases deriving from increased levels of post-mortem damage deriving from ss-DNA library protocol in our newly reported samples, we recalibrated base qualities at transition sites with BaSSdrop (https://github.com/pontussk/baSSdrop) using a SNP list comprising phased (INFO/PHASED=1) transition sites present in autosomes and chromosome X at minor allele frequency (maf)>0.01% in the interim phased data of >200,000 individuals from the UK Biobank Whole-Genome Sequence reference panel (Data-Field 20279). BaSSdrop takes advantage of the fact that sequencing reads generated from ssDNA libraries retain strand-orientation^112^: for transition sites, baSSdrop rescales the base qualities to zero of all cytosines and thymines in forward-oriented reads and all guanines and adenines in the reverse-oriented reads. BaSSdrop does the base quality rescaling independently of reference sequence, focussing instead on transition sites with low maf, to avoid introducing reference bias. We confirmed that the effect of post-mortem damage was corrected by projecting pseudo-halpoid genotypes (before and after baSSdrop recalibration) on a PCA and that imputation accuracy was improved (**Extended Data Fig. 1**).

GRCh38-remapped and recalibrated genomes were filtered for mapping quality >30 (MQ30) and autosomes were imputed with GLIMPSE2^14^ using the low-coverage WGS imputation pipeline^16^ and the UKB reference panel^15^ for a total of ∼660M variants (>200,000 individuals; SHAPEIT5 phased data, Data-Field 20279) on the UK Biobank DNAnexus platform (https://ukbiobank.dnanexus.com/). We applied the same imputation pipeline, but without the recalibration step, to a subset of our newly reported samples subjected to double-stranded library preparation (**Extended Data Table 1**), as well as for published genomes.

We validated baSSdrop recalibration and imputation directly, by testing imputation accuracy for different coverages and maf using GLIMPSE2_concordance (*--af-tag RAF --bins 0.01 0.05 0.1 0.25 0.5 --min-tar-gp 0.8 --gt-val*), comparing genotype likelihoods for downsampled genomes with genotypes called directly from the original high-coverage genomes (7.5–10.3x) as the ground truth, following Sousa da Mota et al.^123^ (**Extended Data Fig.1c**, **Supplementary Fig. 8**). We further validated our approach empirically, by comparing the results based on recalibrated and imputed versus non-recalibrated and non-imputed datasets (*e.g.* ancIBD versus KIN results, *qpAdm* models on imputed and non-imputed genotypes) (**Supplementary Fig. 9; Extended Data Table 11, 12**).

### Uniparental haplogroups

We classified Y-chromosome lineages using Yleaf^124^ v.3.1 (*-r3*, *-q30*, *-dh*, *-hc*) and cross-checked against YFull YTree v.11.01.00 (https://www.yfull.com/tree/) and ISOGG Y-DNA Haplogroup Tree 2019-2020 (https://isogg.org/tree/) (**Extended Data Table 1**). Prior to running Yleaf, we applied baSSdrop recalibration to haplogroup-diagnostic transition SNPs used for haplogroup classification (using both the ‘new’ and ‘old’ positions lists provided by Yleaf).

For mtDNA haplogroup classification (**Extended Data Table 1**) we used Haplogrep2^125^ based on PhyloTree^126^ v.17. We calculated frequencies of mtDNA haplogroups for sites with n≥5. For sites with n≥10 we annotated a site as displaying a ‘matrilineal burial practice’ pattern if one or more haplogroups were present at >10% frequency; for sites with 10> n ≥5, we annotated as a ‘possible’ signal of matrilineal burial practice if a given haplogroup was present at least twice and as ‘yes’ if present at ≥50% (**Extended Data Table 3**). These sample sizes denote the number of individuals after excluding close relatives up to 3rd degree. We calculated Simpson’s *D*, Shannon’s *H* and haplogroup (*h*) diversity indices for all Iron Age and Roman-period sites with *n*≥5 (**Extended Data Table 2**).

### Genotype-based population analyses

We called pseudo-haploid genotypes using samtools *mpileup* (*-R,-B,-q30,-Q30*) and pileupCaller with*--randomHaploid* (sequenceTools v.1.5.2; https://github.com/stschiff/sequenceTools), using two different SNP lists: 1) the ‘1240k’ panel^29^ (in this case also with --singleStrandMode for the ssDNA libraries), and 2) ∼3,868,200 biallelic transversions with 1% maf on the 1KGP phase 3 global panel^127^ (as described in Silva at al.^36^), hereafter referred to as ‘1KGP transversion sites’.

We projected newly sequenced ancient from Britain (as pseudo-haploid genotypes) on a Principal Component Analysis (PCA) computed using genotypes of 1217 present-day individuals from various populations from Europe, Near East, Caucasus and North Africa^128–130^, genotyped at ∼600,000 sites from the Human Origins (HO) array using smarpca (*shrinkmode: YES* and *lsqproject: YES*; EIGENSOFT^131^ v.6.1.4).

We modelled each individual using a 3-source *qpAdm* model with a fixed set of outgroups and sources (adapted from Patterson et al^10^; ADMIXTOOLS^130^ v.5.0; https://github.com/pontussk/qpAdm_wrapper)), based on genotype frequencies calculated using the ‘1KGP transversion sites’ (**Extended Data Table 11**). We confirmed the results on a subset of these individuals using imputed genotypes and Twigstats (more details below).

### Relatedness inference

We inferred biological relatedness with both KIN^132^ and ancIBD^27^ v.0.6 (**Extended Data Table 7**). We used KIN to identify pairs of individuals related up to third degree within sites with a sample size of at least five individuals, using 1KGP transversion sites. We pooled individuals from sites with *n*<5, stratifying by broad archeological period (Bronze Age, Iron Age, Roman period, Early Medieval), to avoid potential biases due to population structure.

We merged the data for individuals identified as identical (with samtools^119^ v.1.3.1 *merge*), and retained them in the dataset as a single individual. We included all individuals in analyses performed independently for each individual (*e.g.* PCA, hapROH, *qpAdm* when used as targets), but excluded individuals related up to third degree for analyses at the population level (e.g. *qpAdm* source populations, Relate/Twigstats, selection scans) by retaining only the individual with the highest coverage.

For more distant relatives, and to identify relatives across different sites and regions, we inferred segments Identical-By-Descent (IBD) shared between pairs of individuals, on a dataset comprising genomes from Britain (*n*=1516, 983 of which newly reported in this study) and neighbouring regions across Eurasia and the Mediterranean (*n*=1161, all previously published) (**Extended Data Table 6**). We used ancIBD, restricting the analyses to imputed genomes with autosomal coverage >0.3x (for shotgun genomes) or >600,000 SNPs (for capture data), resulting in a total of 2677 individuals. We restricted the subsequent IBD analysis to sites overlapping with the ‘1240k’ panel, as in the default mode of ancIBD (making use of our large dataset to calculate allele frequencies), and using a lifted over version of the variants reported in the AADR generated with Picard (http://broadinstitute.github.io/picard/) v.2.23.8 *LiftoverVcf* tool using the UCSC chain file *hg19ToHg38.over.chain* (resulting in a total of 1,150,106 autosomal SNPs).

We inferred relatedness up to sixth degree by combining the number and total length of IBD segments (requiring at least two IBD-segments >20cM shared between individuals), following Ringbauer et al^27^. To validate the IBD results using the 1240k sites lifted-over to GRCh38 coordinates we compared the ancIBD and KIN outputs for all closely-related pairs (up to 3rd degree): 1) ancIBD identifies the same pairs as KIN, with similar within-degree classification; and 2) we see a high correlation (*r^2^*=0.96) between the total length of IBD1 >12cM inferred with both ancIBD and KIN (assuming the sex-averaged mean recombination rate of ∼1.2 cM/Mb to convert KIN estimates to cM) (**Supplementary Fig. 9**).

We investigated patterns of IBD-sharing within Britain and between individuals in Britain and other European regions using different IBD length thresholds. Maps were generated using the *rnaturalearth* package (https://github.com/ropensci/rnaturalearth) on R v.4.4.1. We excluded related individuals up to 3rd degree from the calculations of average pairwise IBD sharing within sites.

### Homozygosity scan

We inferred runs of homozygosity (ROH) with hapROH^99^ (with default parameters and pseudo-haploid mode) on 885 newly generated genomes and 602 previously published individuals from Britain passing the 400,000 SNP threshold, following hapROH recommendations (**Extended Data Table 5**).

### Twigstats framework

We estimated genome-wide genealogies on imputed genotypes with Relate^34^. First, we used the setGT plugin on *bcftools* v.1.19 to set all sites with genotype probability (GP) <80% as missing and exclude sites with missingness >2% (*bcftools +setGT ---t q-n.-i’SMPL_MAX(FORMAT/GP)<=0.8’ | bcftools view-i’F_MISSING>0.02’*). To avoid any additional potential biases caused by postmortem damage and differential treatments across different studies, we further restricted the dataset to transversion sites only. We converted sample dates to generations assuming 28 years per generation^133^ and using the average date in BP (if directly radiocarbon dated) or the midpoint of a range (for contextual dates), following the ADDR notation for consistency between our data and published datasets. To account for a mismatch between present-day genomes, collected in the last ∼20 years from adult individuals, and ancient genomes with dates in BP (which are relative to the year 1950 CE), we added 50 years to all ancient dates before converting years to generations. We used the GRCh38-specific strict mask and ancestral reference files to prepare the input files (with *RelateFileFormats*) and ran Relate with the default transversions-only mutation rate (4e-9), using the autosomal coalescence rates estimated from the 1000 Genomes (input files provided in https://zenodo.org/records/15179498).

We used Twigstats^21^ to calculate time-stratified *f*2-blocks (applying the function *f2_blocks_from_Relate*) on R v.4.4.0-gfbf-2023b, using the GRCh38 recombination map (provided in https://zenodo.org/records/15179498). We calculated *f*2-blocks with two different generation cutoffs: 1) we used a cutoff of 5000 generations (*t*=5000), to capture more ancient variation; and 2) a cutoff of 1000 generations (*t*=1000), to capture more recently shared variation for a focus on post-Bronze Age Europe. We computed pairwise outgroup-*f*3 (with the formulation *f3(Yana; Ind1, Ind2))* from the *f2* blocks (*t*=1000). We stored the results as 1-*f3(Yana; Ind1, Ind2)* genetic distances in a symmetric matrix that was used for MDS and clustering analyses, following the approach by Speidel et al^21^.

### *qpAdm* formulations

We applied different *qpAdm* formulations, using the Twigstats framework, with different combinations of distantly and more proximally temporal sources. We first modelled distal ancestries for all newly reported individuals with a 3-source model with fixed sources (*t*=5000, using Mesolithic, Neolithic and Early Bronze Age individuals as reference populations^10^) (**Extended Data Table 12**).

For individuals in the last 2000 years we used *qpadm_multi* to run every combination of 1-, 2-, and 3-source models largely based on a set of references curated to model the ancestry of the targets within the context of European Iron Age and Roman-period groups^21^. We considered a model accepted (i.e. not statistically rejected and plausible) if *p*-value>0.01 and *ancestry proportion±SE* between 0 and 1. We prioritised lower-ranking models, applying the following rationale for each *n-*ranking model: 1) for each individual target, if the *n*-ranking model was accepted, we did not explore higher-ranking models; 2) if no *n*-ranking model was accepted we considered *n+1*-ranking models. In the cases of multiple equally complex models being accepted we plot only the model with the highest *p*-value, but report all results in **Extended Data Tables 8 and 9**. We applied tailored *qpAdm* formulations as follows:

***Iron Age.*** We pooled Iron Age individuals dating to post-200 BCE in two groups as targets and rotating through combinations of 1, 2 and 3 Iron Age sources (from France and Scandinavia).

***Roman.*** We applied an additional model including also the late Iron Age individuals from southeast Britain as a possible source (*Iron Age Britain post-200 BCE*).

***1st millennium CE.*** We merged the two Scandinavia Iron Age groups into a single source. In addition, in the absence of continental genomes with whole-genome data available that could be used as proximal sources, we added a new source population (labelled here’Early Medieval Britain I’) comprising the earliest newly-sequenced individuals from Britain who showed similar ancestry profile in an ADMIXTURE clustering analysis (all dating to the 6th CE), as a proxy source for newly-arrived ancestry from the North Sea coast.

### Clustering: NNLS chromosome painting and ADMIXTURE

We applied Twigstats chromosome painting functions on Relate genealogies, with a block size of 0.05 cM (*blgsize* = 5e-4; *blocksize*=5000), to paint every genome from Britain using a selection of preceding and/or contemporary individual genomes as the palette (**Extended Data Table 13**). We then computed admixture proportions using non-negative least squares (NNLS) on painting profiles.

To maximise the number of individuals we also ran ADMIXTURE on a larger imputed dataset restricting the analysis to 1240k transversions sites. For this, we set all sites with genotype probability (GP) <80% as missing and exclude sites with missingness >10% (*bcftools +setGT ---t q-n.-i’SMPL_MAX(FORMAT/GP)<=0.8’ | bcftools view-i’F_MISSING>0.10’*), and retaining 63,707 LD-pruned biallelic transversion sites, with a maf 1% using PLINK^134^ v.1.9b-foss-2016b (*plink --maf 0.01 --indep pairwise 1000, 50, 25*).

### Single-locus selection scan

We set imputed sites with genotype probability (GP) <80% as missing (with *bcftools* v.1.19 setGT plugin) and excluded sites with missingness >10%. We then identified and excluded individuals with uncertain dates, as well as ancestry outliers based on PC coordinates on a PCA of imputed diploid genotypes (**Extended Data Fig. 5**; based on ∼600k SNPs typed on the HO array, using *smartpca* without projection), including also modern individuals genotyped in the HO array (same as above).

We built two datasets including only imputed ancient genomes from Britain dating from 4500 BP to 800 BP, after excluding outliers: 1) 1257 ancient genomes from Britain, retaining only 1240k sites to maximise the number of genomes, with and combining both newly-generated and previously published data (both shotgun and capture; 806,202 sites retained after filtering for missingness <10% and maf >0.05); and 2) 766 individuals, comprising only our newly-sequenced whole genomes, for a total of 5,562,421sites (missingness <10% and maf >0.05). For ancestry-restricted selection scans, we additionally restricted the individuals on both dataset 2, by retaining only individuals dating to the Bronze and Iron Ages (after outlier removal), plus Roman-period and early medieval individuals modelled as deriving ancestry entirely from Britain Iron Age (based on Twigstats-boosted *qpAdm* modelling), resulting in a total of 427 individuals retained.

We ran a linear regression with PLINK using time in years BP as phenotype on both datasets^86^, and applied standard genomic correction to adjust *p*-values^29,86,135^ and confirmed the results applying a Genetic Relationship Matrix (GRM) and a Mixed Linear Model (MLM) approach with GCTA^136^ (*--mlma--grm*). All *p*-values reported in the main text result from the PLINK linear regression. Linkage disequilibrium between SNPs was estimated using LDlink (https://ldlink.nih.gov/ldpair) and allelic geographic distributions were plotted with GGV (https://popgen.uchicago.edu/ggv).

## Extended Data Figures

**Extended Data Fig. 1:**
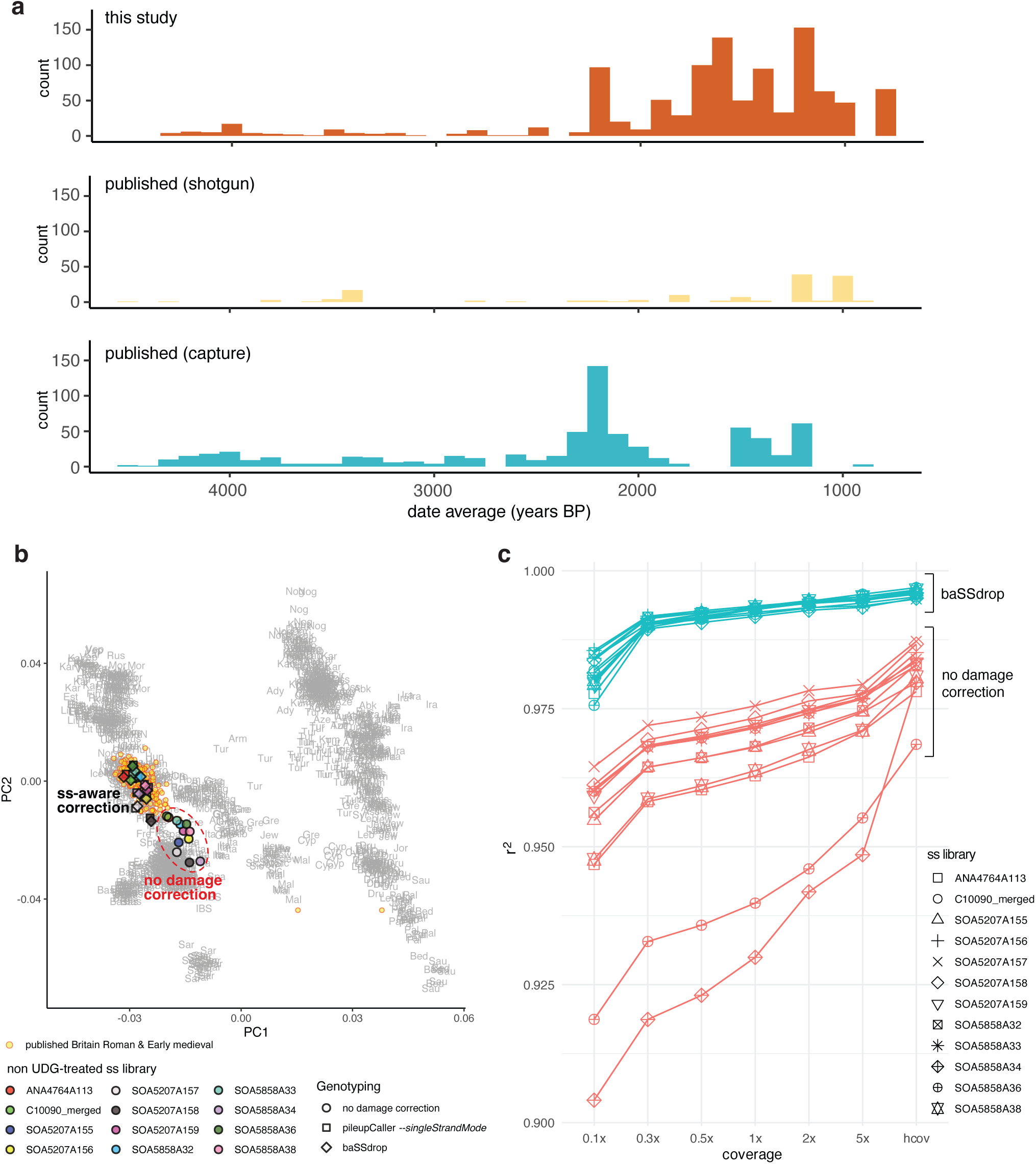
Time transect of whole shotgun genomes from Britain. **a)** Comparison of the dataset generated in this study with previously published aDNA data from Britain with metadata reported in the AADR v.62 (both whole-genome shotgun and capture data), focussing on individuals with average dates between 4500 and 800 years BP, in 100-year bins. **b)** Principal component analysis (PCA) with aDNA ss libraries projected on PCs computed using present-day western Eurasians genotyped on the Human Origins (HO) Affymetrics panel (∼600k SNPs). Each high-coverage aDNA library was genotyped using three strategies: 1) without applying any damage correction, 2) with pileupCaller using *--singleStrandMode* and 3) recalibrating damage using baSSdrop (**Methods**). **c)** Imputation accuracy plots (*r*^2^) for different downsampled coverages (5x, 2x, 1x, 0.5x, 0.3x and 0.1x), comparing imputation with and without applying baSSdrop damage correction. Initial coverages (hcov) varied 7.5–10.3x after remapping to GRCh38 and filtering for MQ>30 (**Methods**).

**Extended Data Fig. 2:**
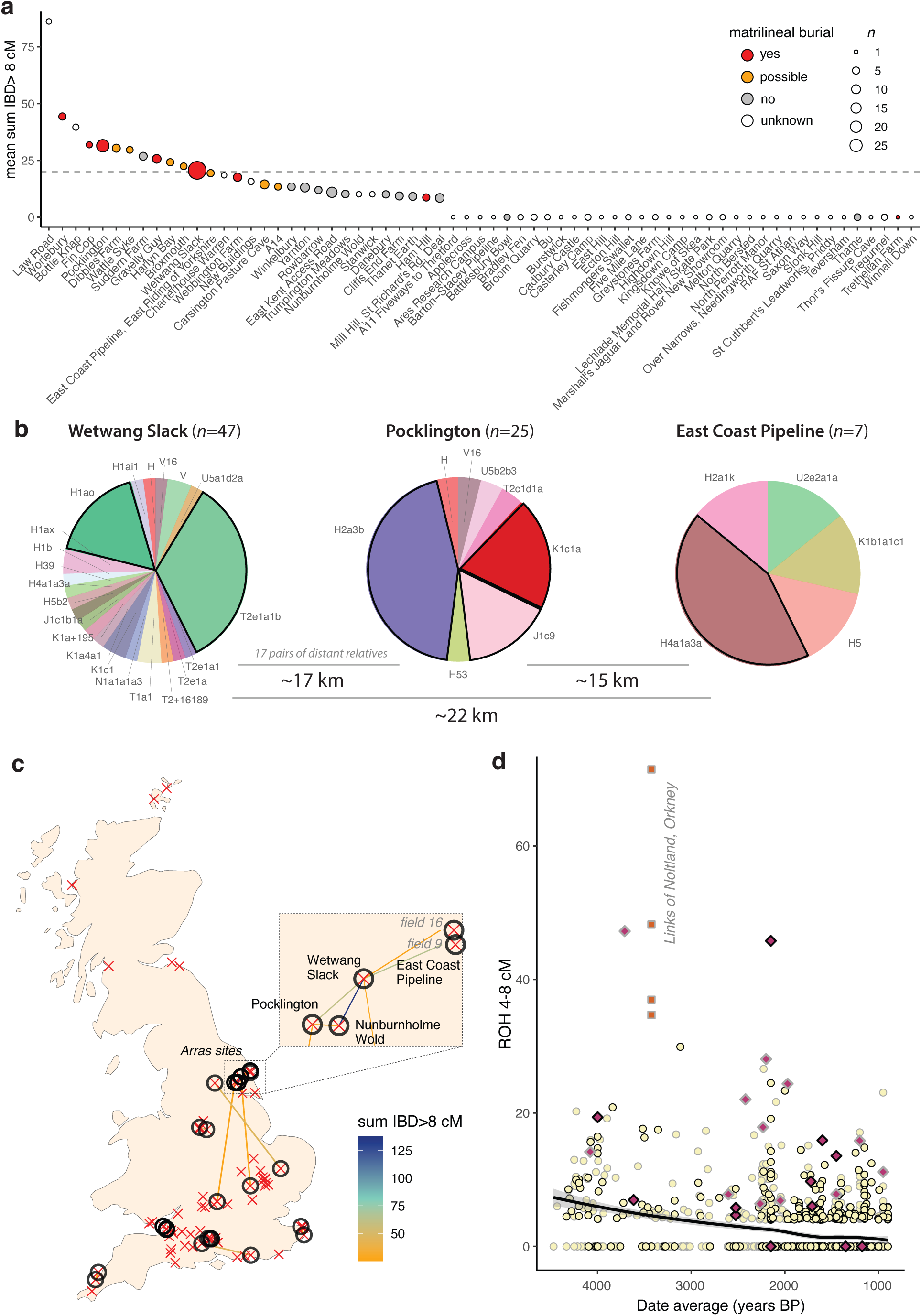
Patterns of relatedness in the Iron Age. **a)** Mitochondrial DNA haplogroup frequency in three Iron Age ‘Arras’ sites in Yorkshire (excluding relatives up to 3rd degree). Haplogroup frequencies for other Iron Age and Roman-period sites in **Supplementary Fig. 3 and 4. b)** IBD sharing between Iron Age sites. Sites with one or more individuals sharing at least one IBD segment >25 cM with another individual in a different site are represented by circles and connected by a line coloured according to the total sum of shared IBD (>8 cM) shared between the pair. Remaining sites included in the analyses are represented by a single red ‘x’. Dataset includes shotgun and capture individuals, excluding relatives up to 3rd degree (**Extended Data Table 6 and 7**). **c)** Decrease in short runs of homozygosity (ROHs; 4-8 cM), suggesting increased effective population sizes over time. The analysis was restricted to individuals with >400K autosomal SNPs overlapping with the 1240k panel; newly-generated genomes with a black outline, previously published data from Britain with grey outline. Local regression (LOESS) excludes individuals with evidence of consanguinity (defined as having a total of >50 cM of the genome in ROHs >12 cM^99^; represented by purple diamonds) and individuals from the site of Links of Noltland, in Orkney, previously reported as sharing increased long ROH, evidencing low population size^76^ (represented by orange squares). **d)** Mean intra-site IBD-sharing in the Iron Age. Data points coloured according to matrilineal burial practices detected in the site, with size reflecting sampling size excluding relatives up to 3rd degree. Dashed line at 20 cM.

**Extended Data Fig. 3:**
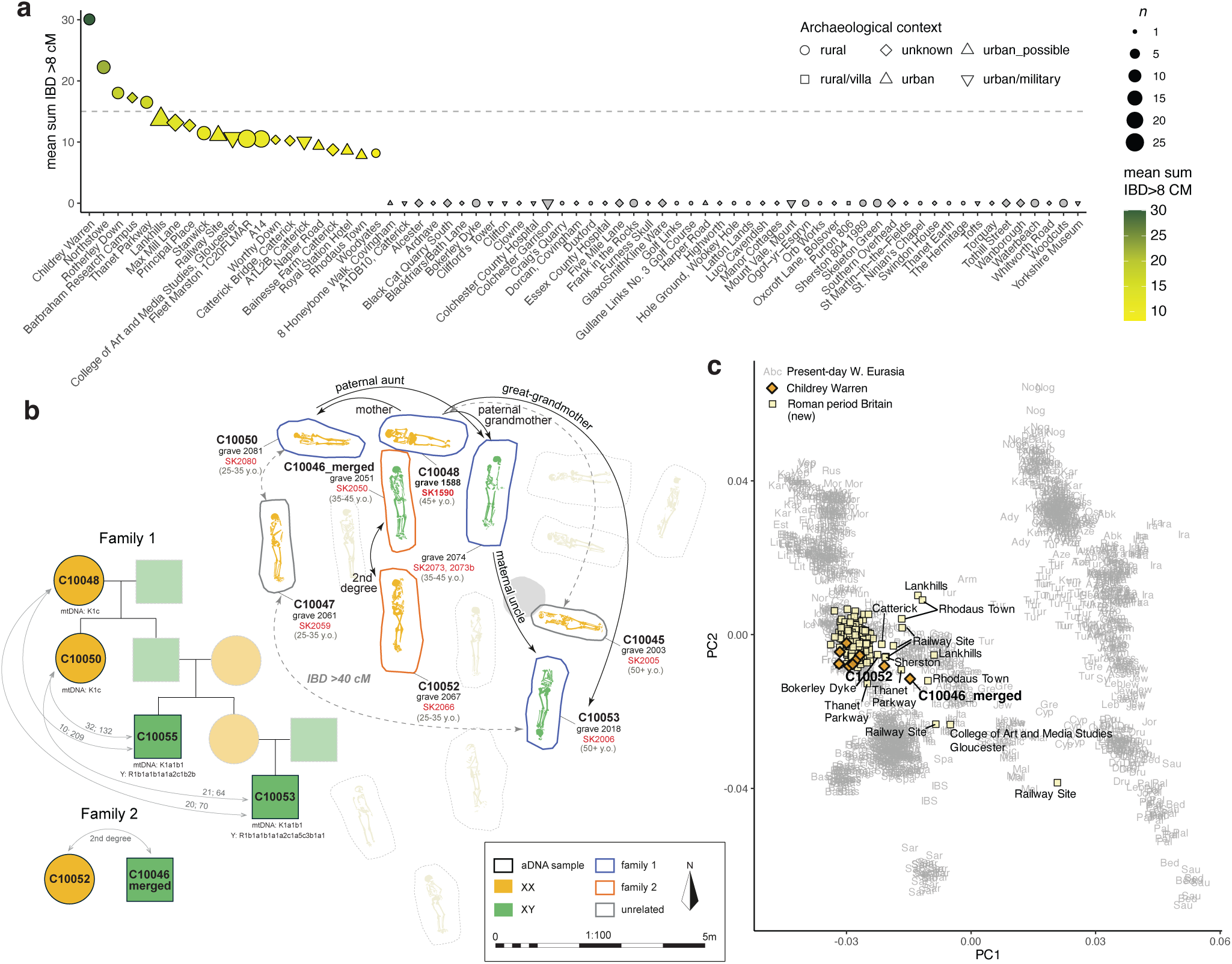
Intra-site IBD-sharing and relatedness in Roman Britain. **a)** Intra-site mean IBD-sharing in the Roman period (only newly-reported sites shown). Data points coloured according to mean sum IBD >8 cM, with shape denoting archaeological context and size reflecting sample size. Dashed line at 15 cM. **b)** Biological relatedness within the Late Roman Childrey Warren cemetery. Reconstructed family trees for two family groups identified within the cemetery and plan of the excavated area with sampled individuals coloured according to karyotypic sex and grave outline according to family group. **c)** PCA projecting all newly-sequenced Roman-period individuals (yellow squares), with Childrey Warren individuals highlighted (orange diamonds) on PCs calculated using present-day individuals from western Eurasia, genotyped on ∼600k Human Origins (HO) sites. Outlier individuals from Childrey Warren are individually labelled; all other outliers are labelled with site name.

**Extended Data Fig. 4:**
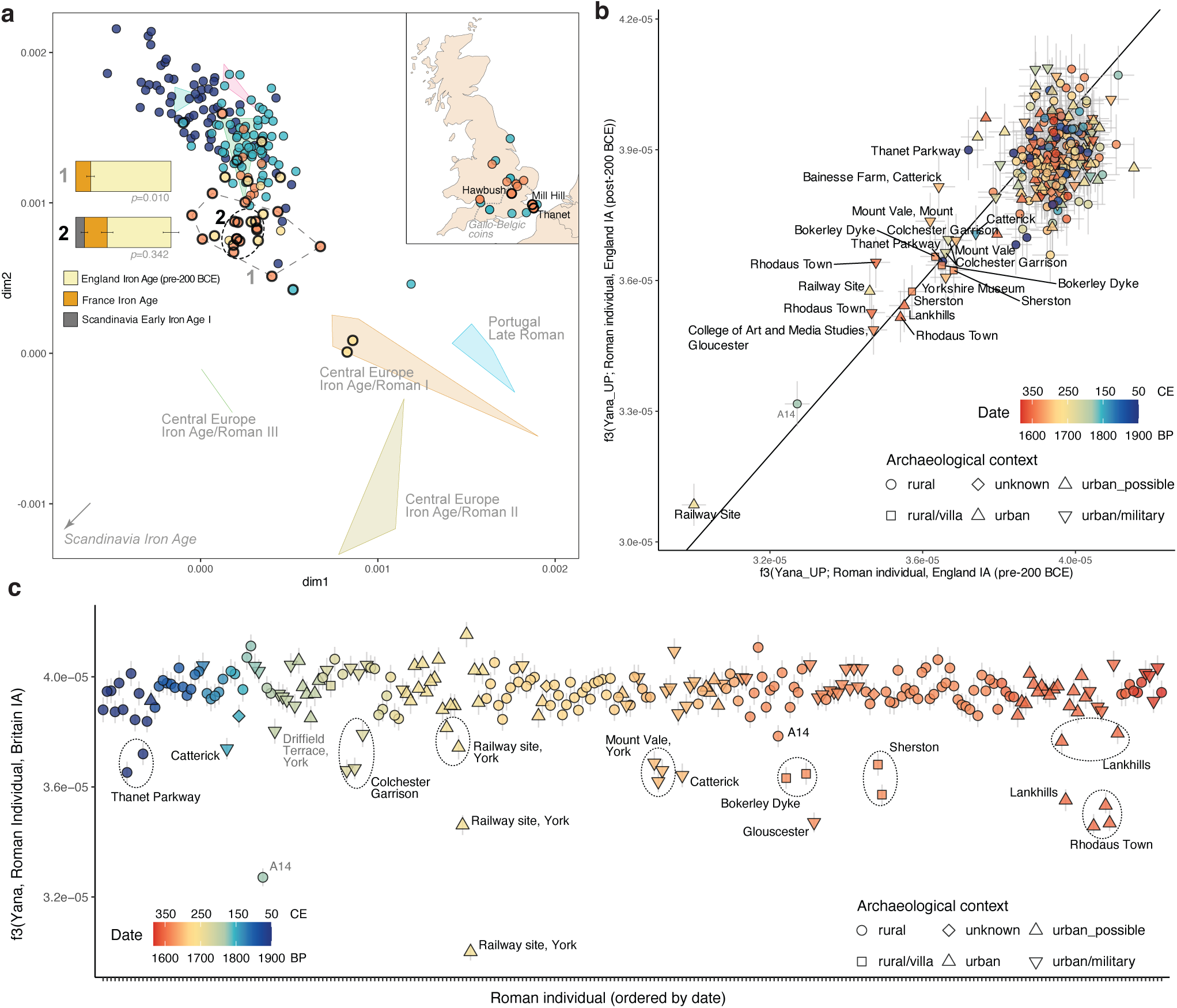
Ancestry in Roman Britain. **a)** MDS based on Twigstats-boosted pairwise *1 - outgroup-f3* estimates (*t*=1000 cutoff). Polygons represent previously published Bronze and Iron Age individuals from continental Europe, Britain and Ireland. Circles represent newly published individuals dating to the Bronze (dark blue), Early-Mid Iron Age (800–200 BCE, light blue), Later Iron Age (200 BCE–0 CE in orange and 0–50CE in yellow). Inset map shows archaeological sites with individuals dated to the Iron Age, with data points coloured as in the MDS and thick borders representing sites where the influx of continental-related ancestry is detected. Ancestry models for Later Iron Age Britain (200 BCE–0 CE) are shown, pooling the individuals inside the dashed circles as targets in two groups. **b)** Comparison of outgroup-*f*3 values using Britain IA (pre-200 BCE) and Britain IA (post-200 BCE). Error bars denote 1 SE. **c)** Outgroup-*f*3 values using Britain IA (pre-200 BCE). In both b) and c) target Roman-period individuals are coloured according to average date in BP and with shape denoting archaeological context.

**Extended Data Fig. 5:**
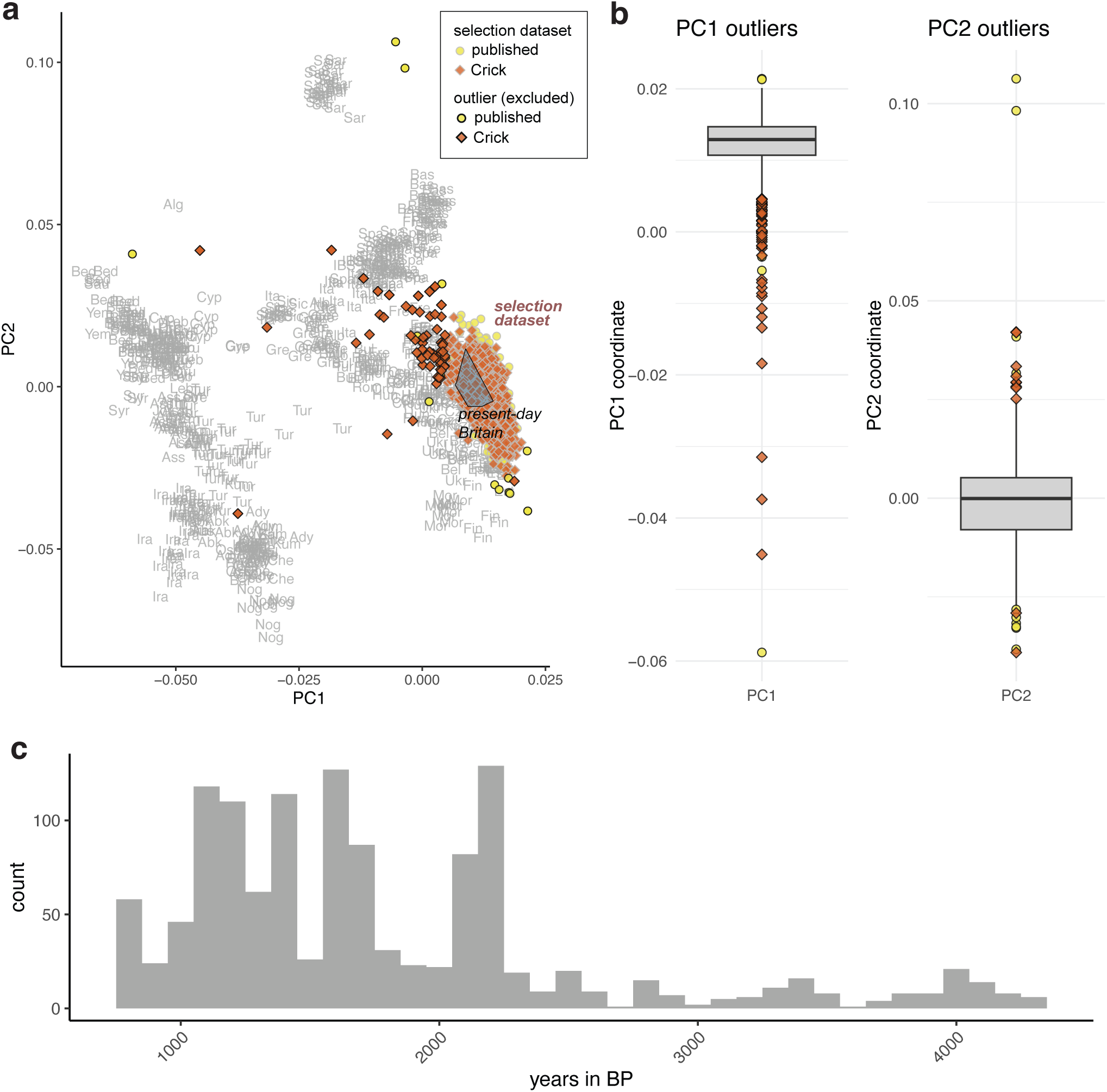
Single-locus selection dataset. **a)** PCA of imputed diploid genotypes (based on ∼600k SNPs), including modern individuals genotyped in the HO array, from Europe, Near East and Caucasus, and ancient genomes from Britain dating from 4500 BP to 800 BP. Present-day individuals from England and Orkney highlighted with a polygon and labelled ‘present-day Britain’ (10 individuals from England and 13 from Orkney, Scotland). Final imputation dataset represented by datapoints with a grey outline. Outliers, as identified in the boxplots shown in panel b, are highlighted with a black outline and were excluded from all the selection datasets. No batch effect is observed between our in-house generated data, labelled ‘Crick’ (non-UDG-treated single-stranded libraries, whole-genome sequencing), and previously published genomes (a mixture of UDG-treated and non-treated, single-and double-stranded libraries, and whole-genome and 1240k-capture data). ‘Crick’ (orange diamonds) includes four previously published genomes^36,100^ that were subjected to the same lab protocols as the newly-published genomes. **b)** Outlier identification based on principal component (PC) coordinates. **c)** Histogram of individuals included in the selection dataset, per 100-years bins.

**Extended Data Fig. 6:**
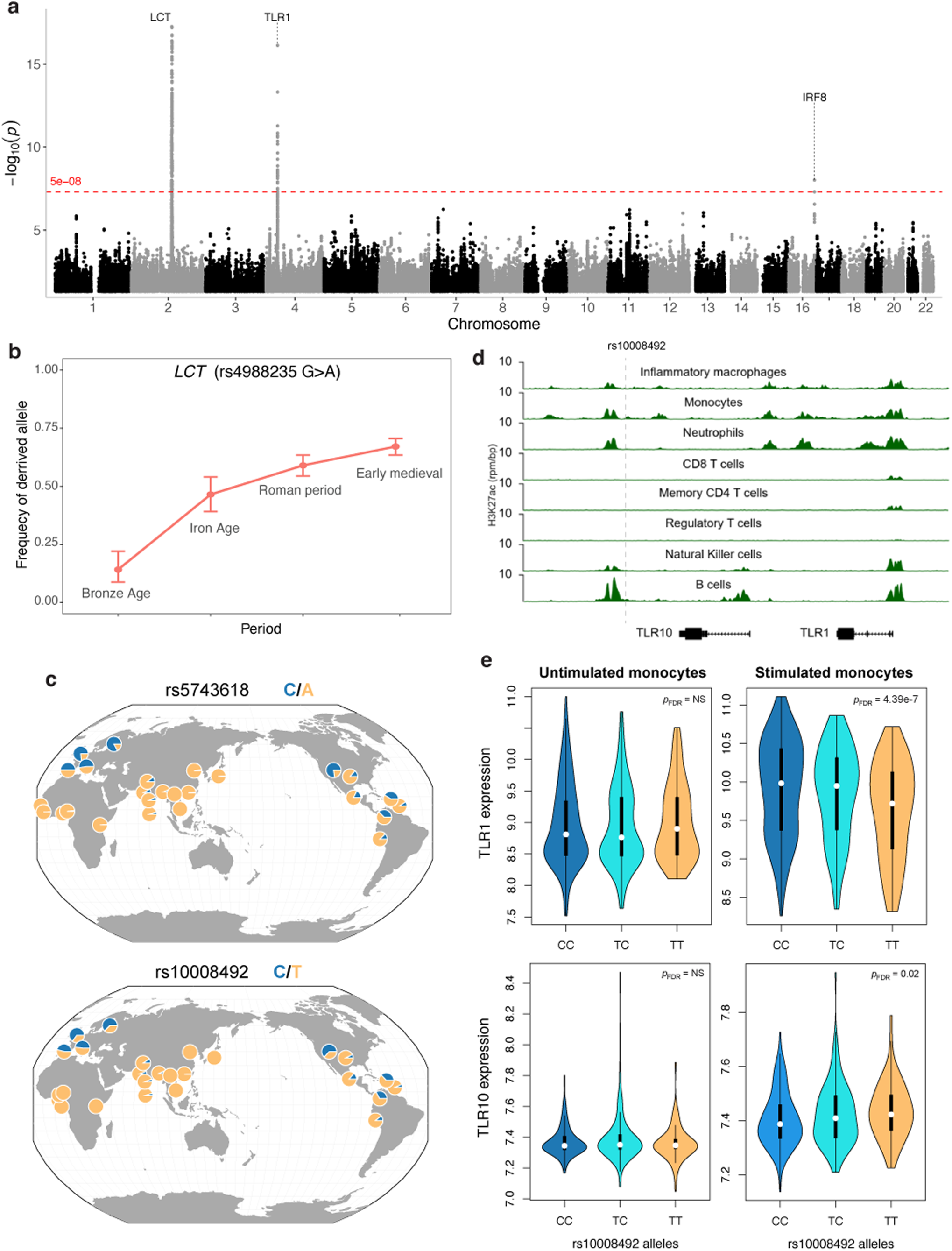
a) Manhattan plot of *p*-values resulting from a mixed linear model using a Genetic Relationship Matrix (GRM) to control for population structure on the ancestry-restricted dataset (*n*=427). **b)** Estimated frequency of the derived allele of rs4988235 on *LCT* in Britain in different archaeological periods. **c)** Global allele frequencies in the 1000 Genomes Project populations of rs5743618 and rs10008492. **d)** Regional variation near TLR10 and TLR1 shown together with H3K27ac tracks for primary immune cells. **e)** Tests for association with the genotype at rs10008492 and expression of TLR1 and TLR10 in stimulated and unstimulated monocytes.

