## Supplementary Fig. for "Genomic history and selection in Roman and early medieval Britain"

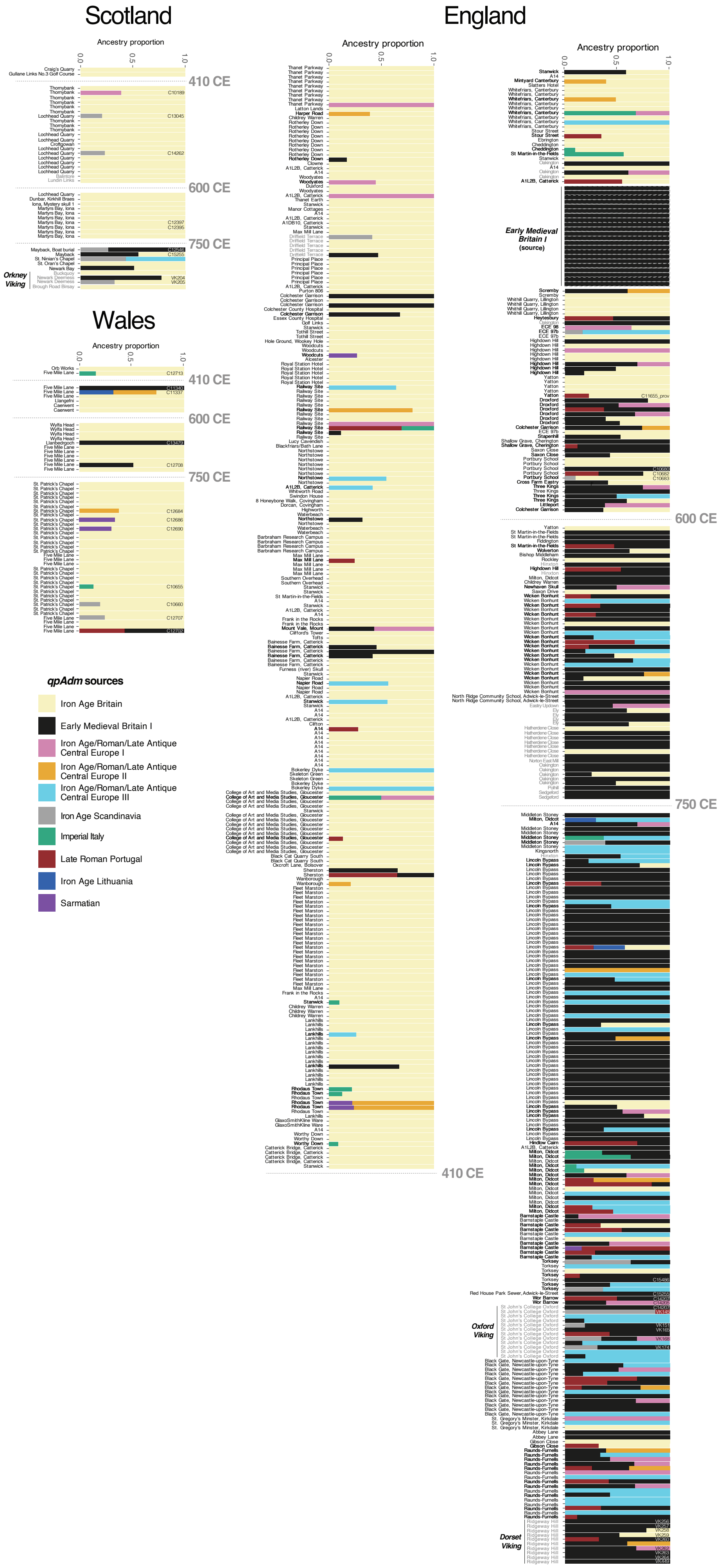
Scotland
England
Wales
Ancestry proportion
0.0
0.5
1.0
410 CE
600 CE
750 CE
Orkney Viking
Early Medieval Britain I (source)
Oxford Viking
Dorset Viking
qpAdm sources
Iron Age Britain
Early Medieval Britain I
Iron Age/Roman/Late Antique Central Europe I
Iron Age/Roman/Late Antique Central Europe II
Iron Age/Roman/Late Antique Central Europe III
Iron Age Scandinavia
Imperial Italy
Late Roman Portugal
Iron Age Lithuania
Sarmatian

**Supplementary Fig. 1:** Twigstats-boosted *qpAdm* models for individuals from Britain dating to the 1st millennium CE (as in Fig. 1d), with site labels for all targets and sample identifiers for individuals highlighted in the main text. Both newly sequenced and previously published (site labels in grey) individuals are included and are plotted according to their average date (Extended Data Table 1); note that the individuals included in the *Early Medieval Britain I* source (all dated to the 6th century CE) are also plotted. Only individuals with accepted *qpAdm* models are shown (*p*-value>0.01; **Methods**). Full *qpAdm* results in **Extended Data Table 8**.

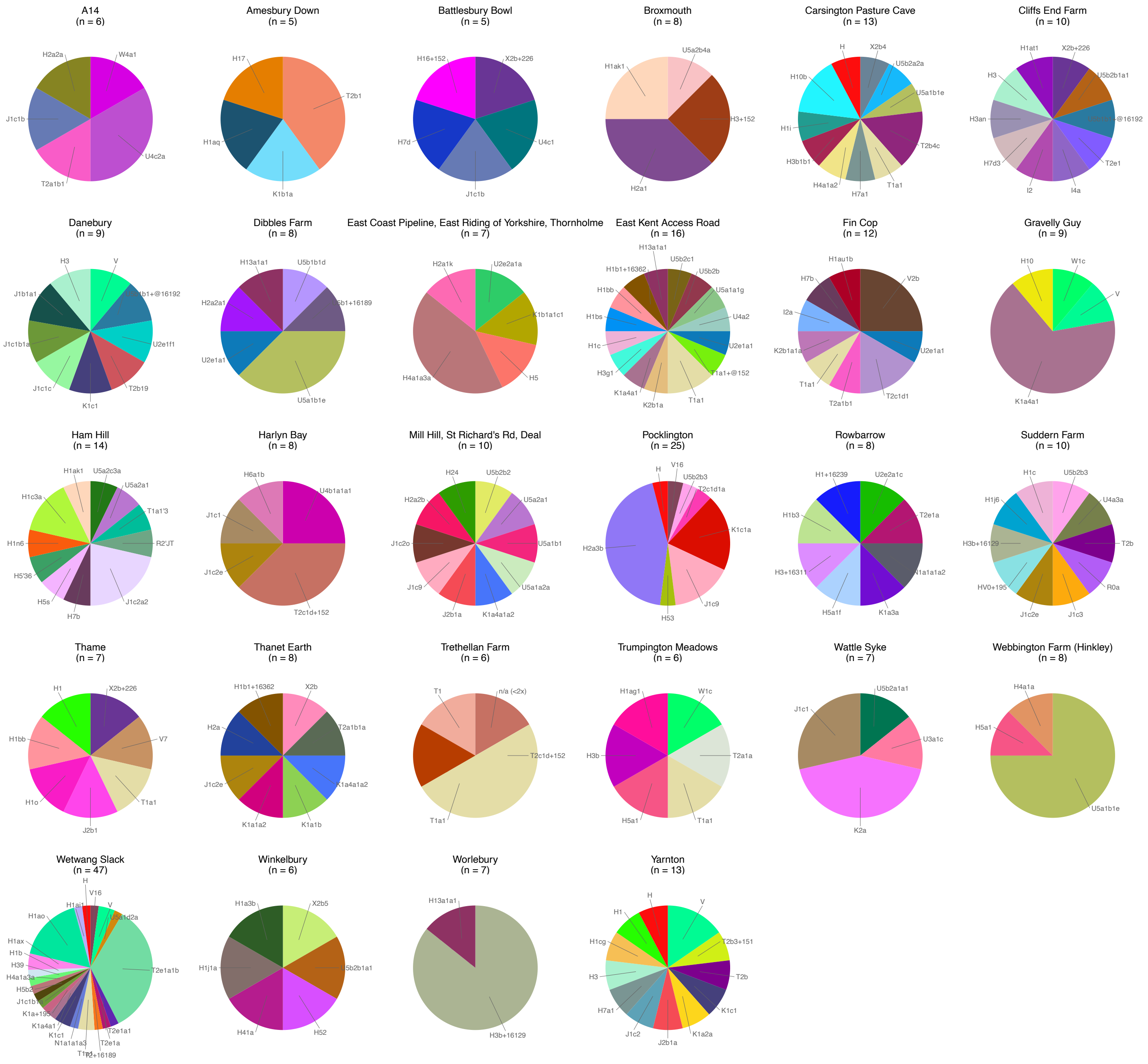


**Supplementary Fig. 2:** Mitochondrial haplogroup frequencies in Iron Age sites with $n \geq 5$ (excluding close relatives).

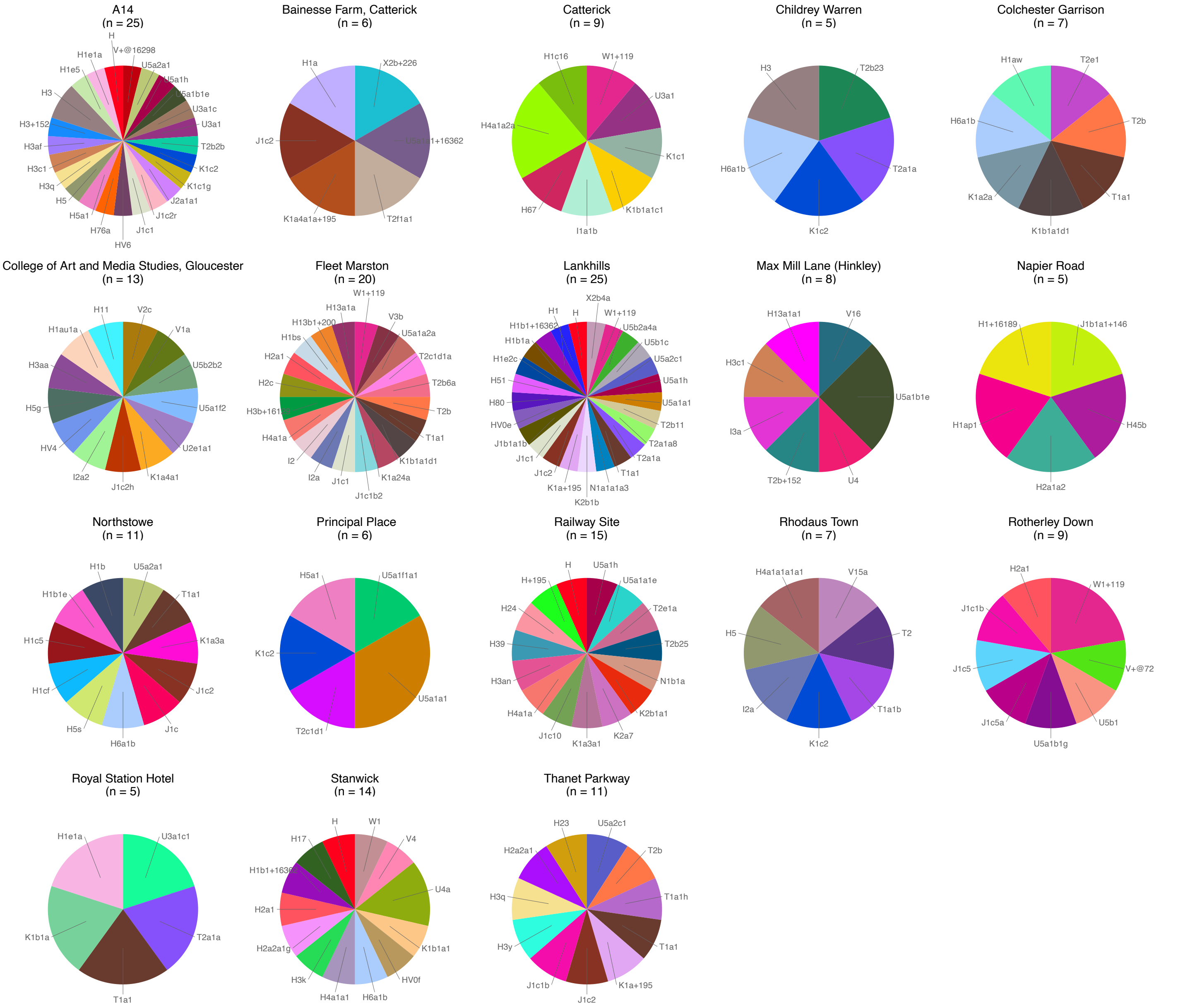


**Supplementary Fig. 3:** Mitochondrial haplogroup frequencies in Roman-period sites with *n*≥5 (excluding close relatives).

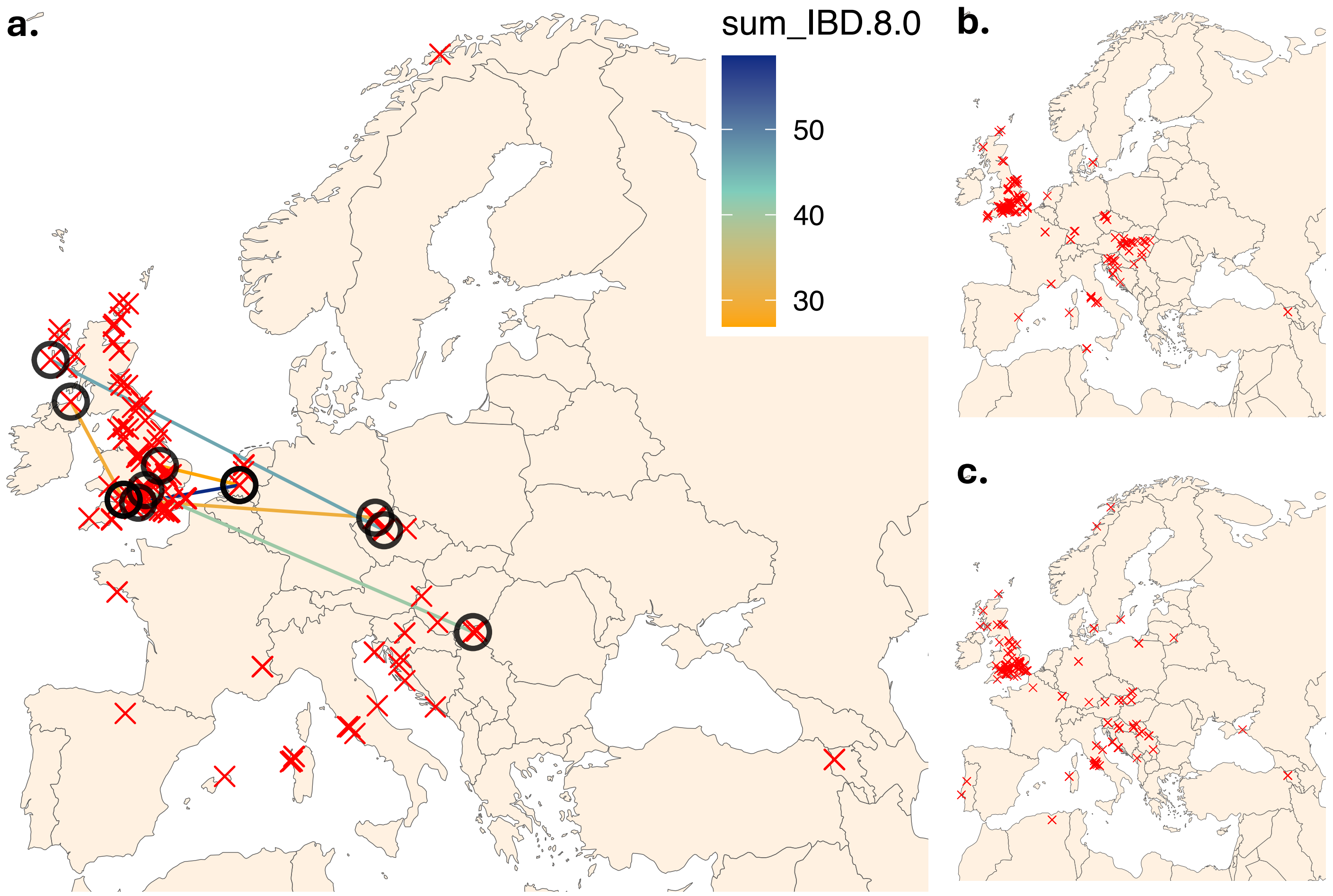


**Supplementary Fig. 4:** IBD-sharing between individuals in Britain and outside of Britain in the **a)** Bronze Age, **b)** Iron Age and **c)** Roman period. Sites with one or more individuals sharing at least one IBD segment >25 cM with another individual in a different site outside of Britain are represented by circles, and both sites are connected and coloured according to total IBD sharing (>8 cM). Remaining sites included in the analyses are represented by a single red 'x'. Dataset includes shotgun and capture genomes (**Extended Data Table 6**). Note that these maps do not show IBD-sharing within Britain or between pairs that do not include one individual from Britain.

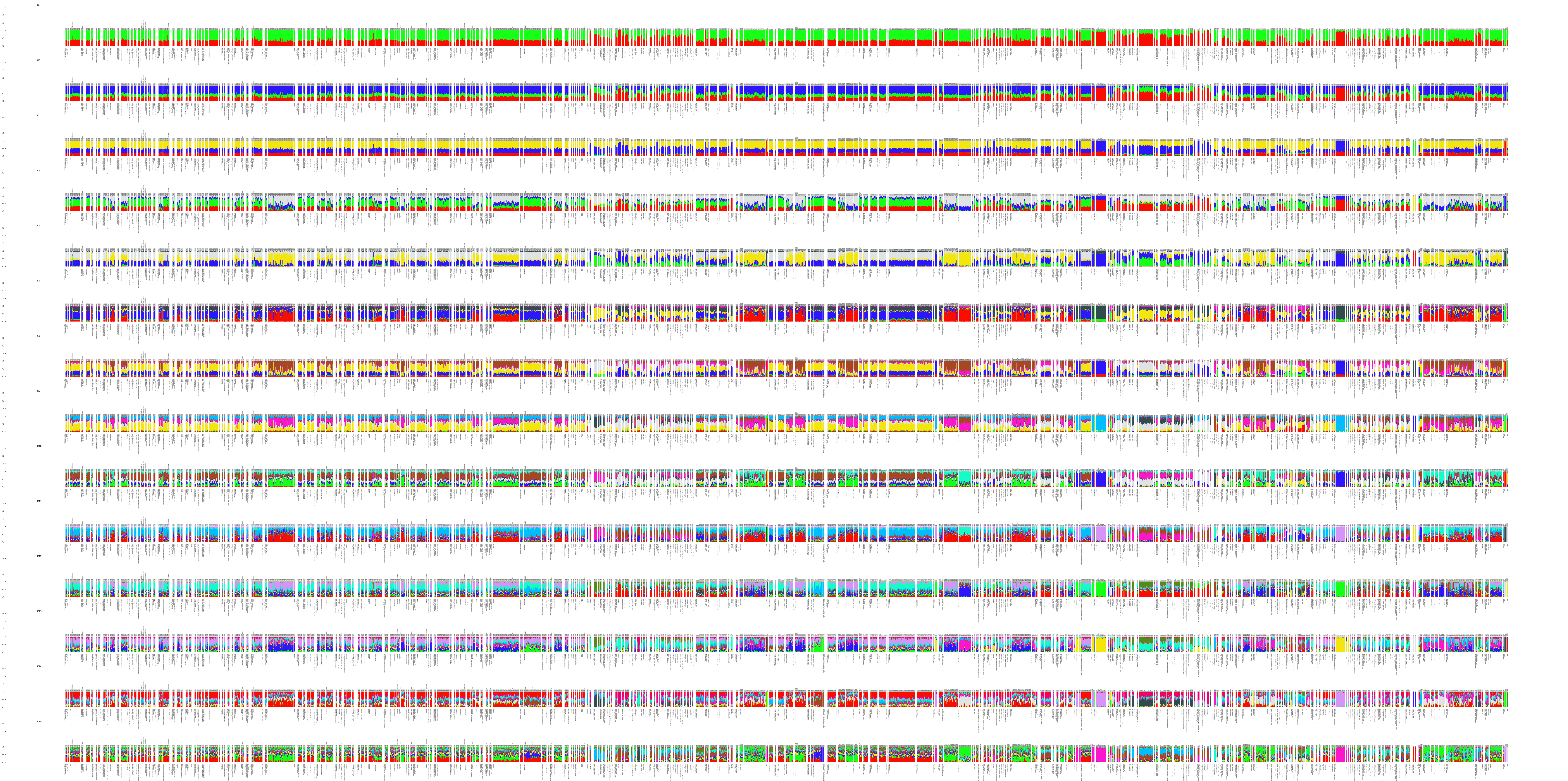
K2
K3
K4
K5
K6
K7
K8
K9
K10
K11
K12
K13
K14
K15

**Supplementary Fig. 5:** Unsupervised ADMIXTURE clustering, from *K*=2 to *K*=15.

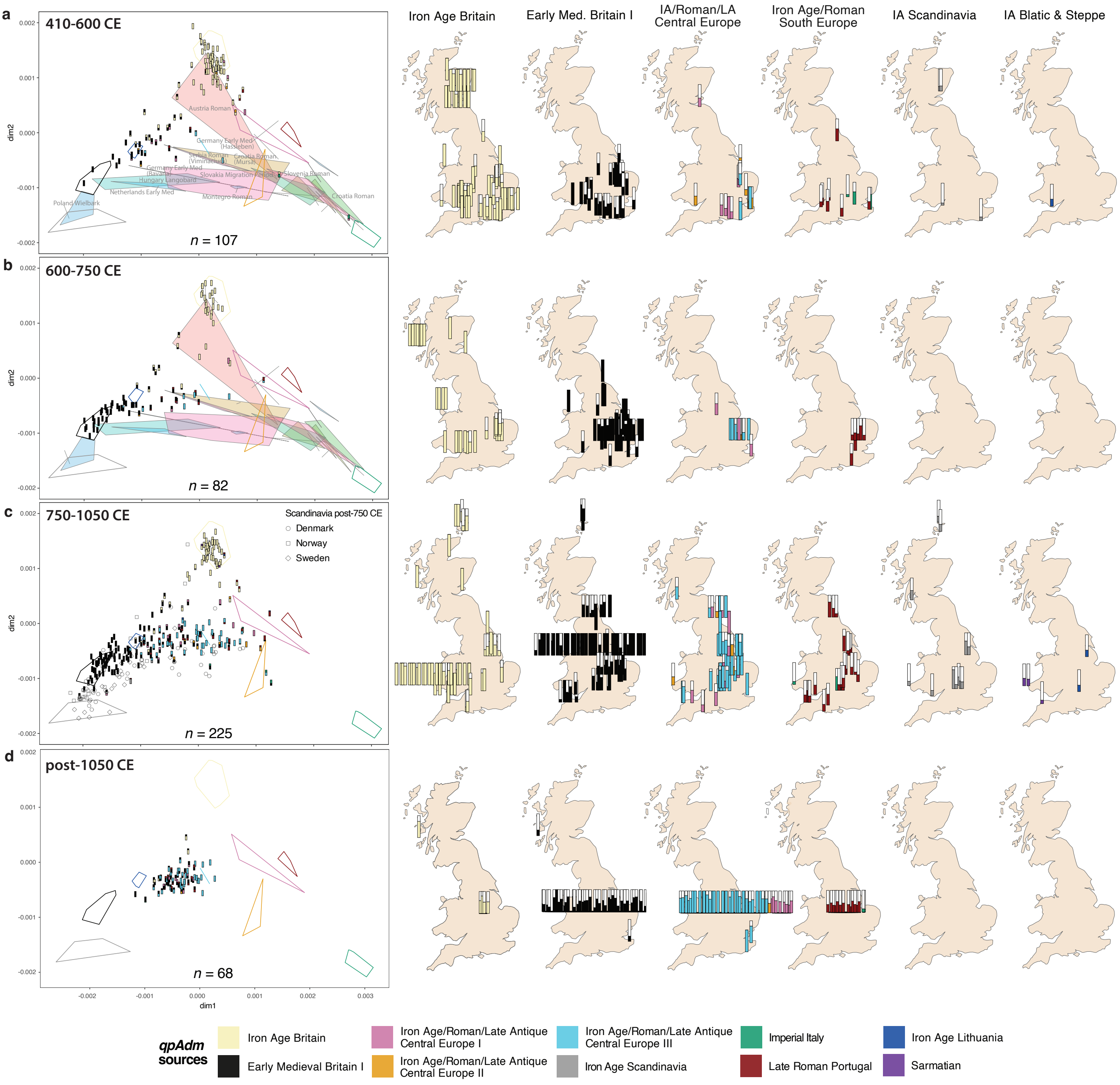


**Supplementary Fig. 6:** Twigstats-boosted MDS and qpADM modelling for early medieval and later targets in different periods: **a)** 410-600 CE, **b)** 600-750 CE, **c)** 750-1066 CE **d)** post-1950 CE. MDS based on Twigstats-boosted pairwise outgroup-*f*3s (*t*=1000 cutoff), stratified by time. Individuals from Britain (both newly sequenced and published) are used as targets and represented by their *qpAdm* modelling proportions (as in Fig. 2; showing only plausible and non-rejected models, i.e. *p*-value>0.01 and ancestry proportion±SE between 0 and 1). Number of targets (*n*) for each time period indicated on each MDS. Polygons represent source populations included in the *qpAdm* models (Sarmatian not included in the MDS) and previously published continental populations dating to the Roman and early medieval periods. Maps show the proportion of ancestry derived from each tested source for each individual. Model selection detailed in **Methods**. All *qpAdm* results in **Extended Data Table 8**.

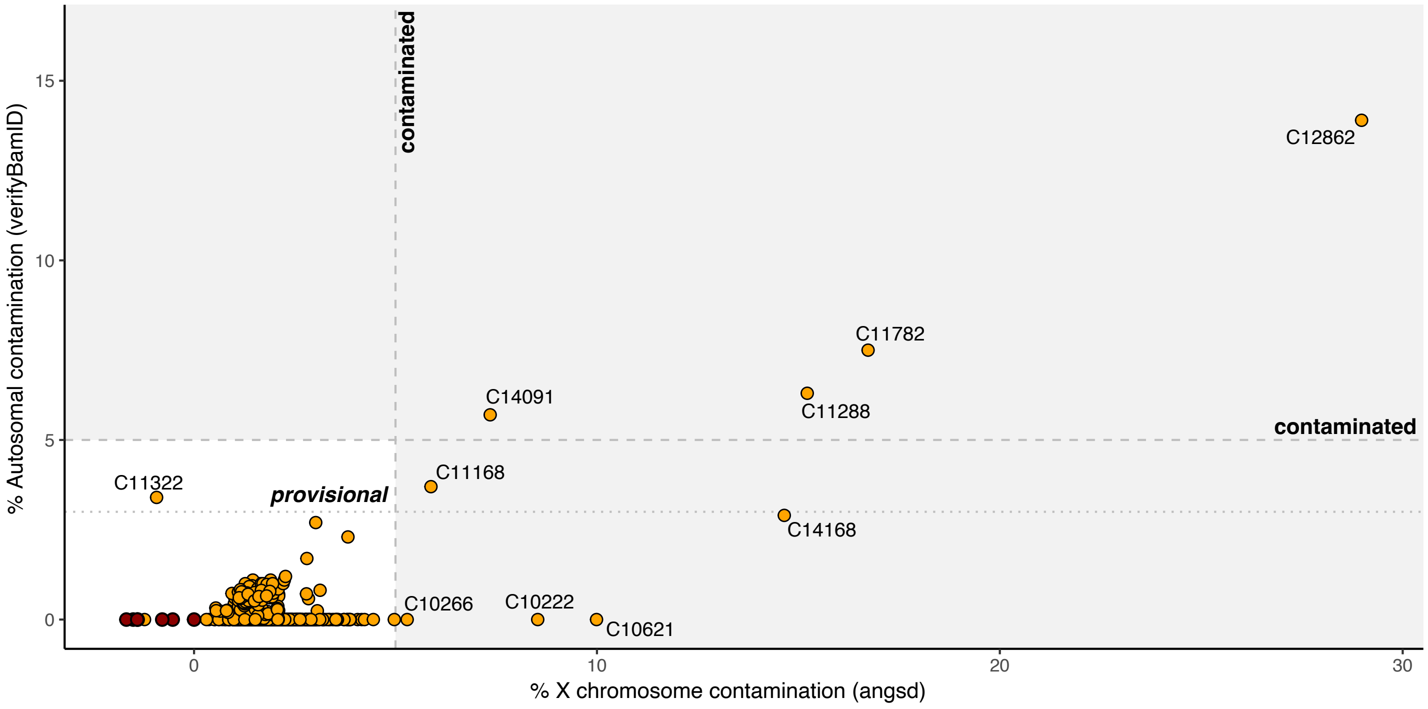


**Supplementary Fig. 7:** Comparison of autosomal (verifyBamID[1]) and X-chromosome (ANGSD[2]) contamination estimated for libraries with a XY karyotype (**Methods**). Individuals with average autosomal coverage (MQ>30) under 0.1x coloured in dark red.

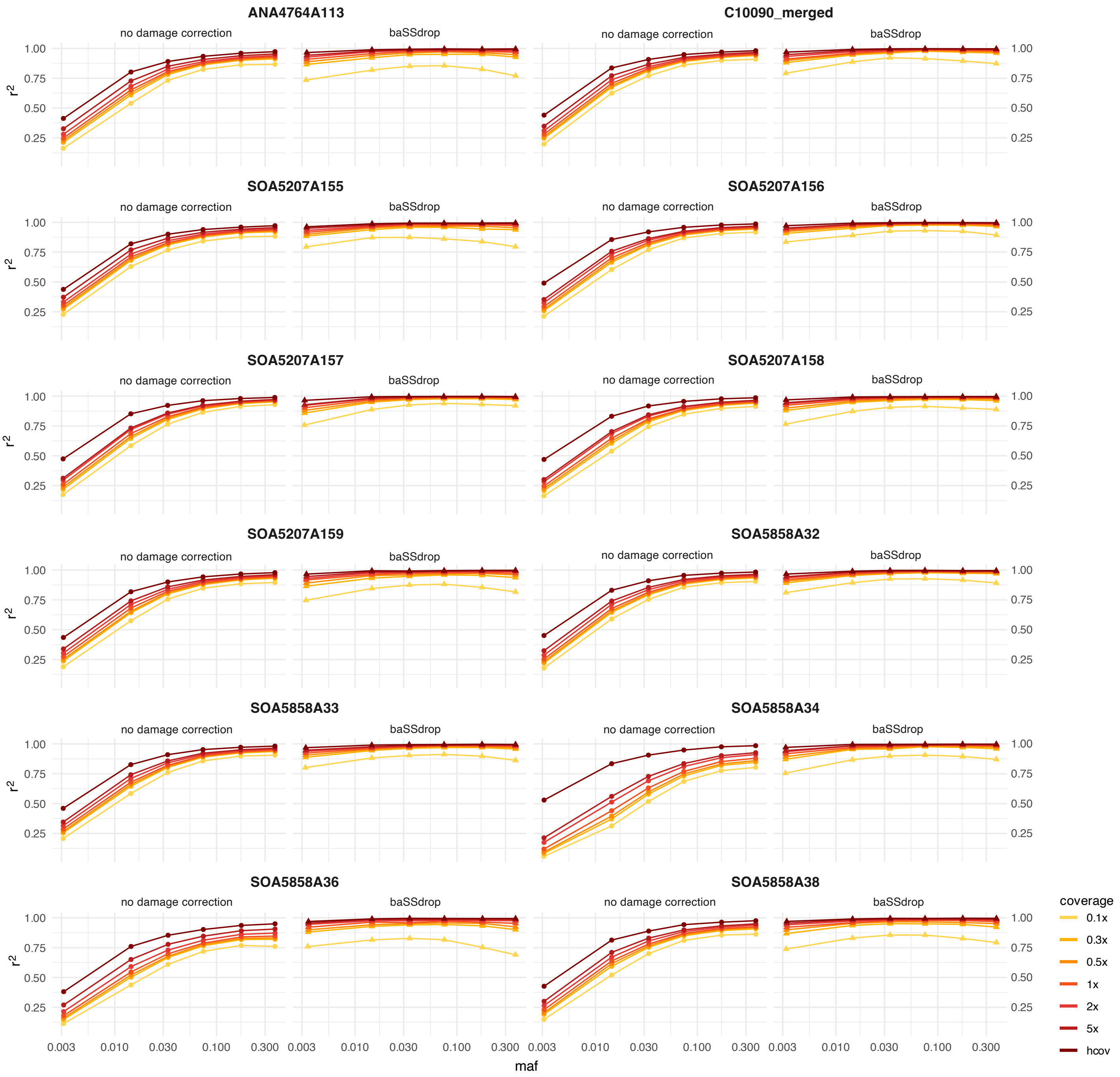


**Supplementary Fig. 8:** Imputation accuracy ($r^2$) for different downsampled coverages (5x, 2x, 1x, 0.5x, 0.3x and 0.1x) at different maf, comparing imputation with and without applying baSSdrop damage correction for 12 high-coverage samples generated using ssDNA library preparation (**Methods**). Initial coverages (hcov) varied 7.5–10.3x after remapping to GRCh38 and retaining only reads with MQ>30.

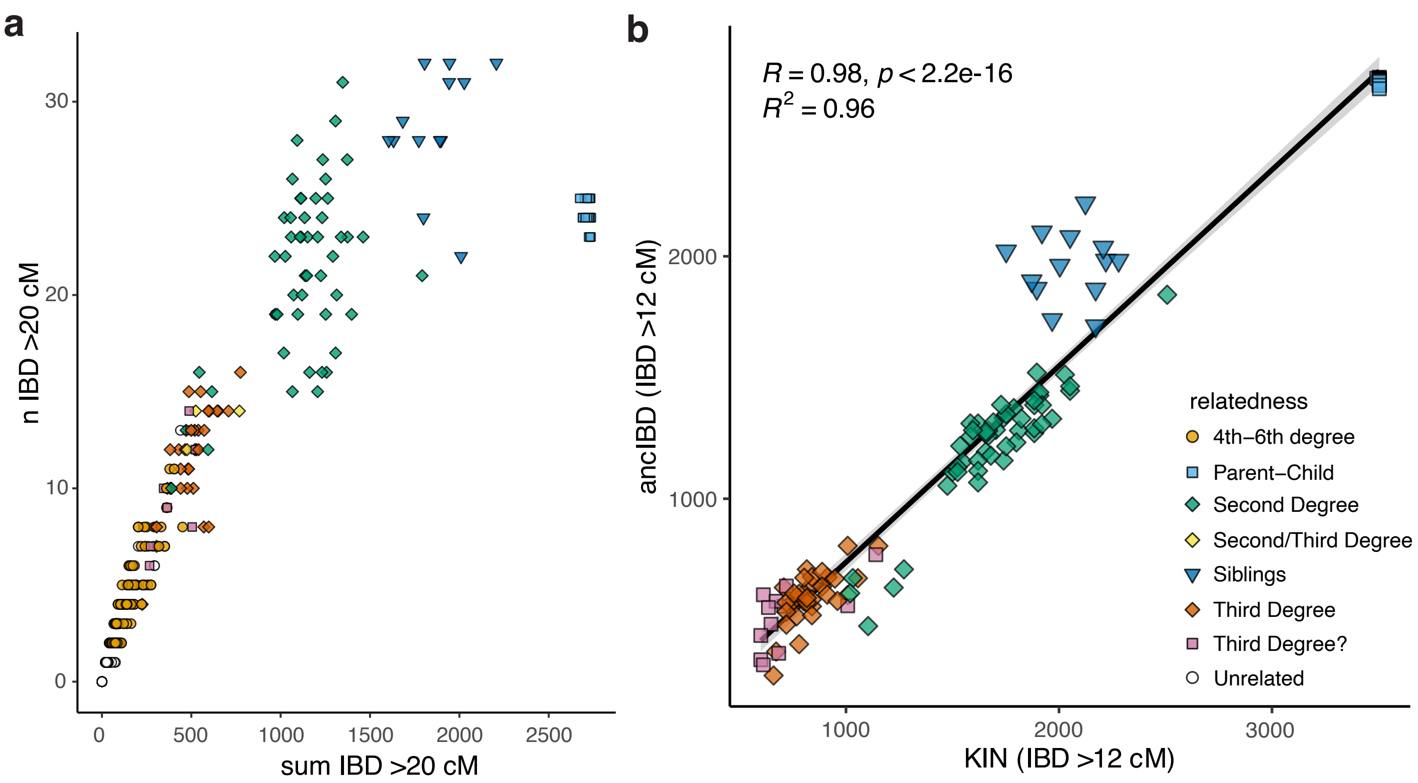


**Supplementary Fig. 9: a)** Sum of IBD1 >20 cM segments and total length of genome in IBD1 (>20 cM). **b)** Correlation in total length of IBD1 >12 cM identified with ancIBD and KIN (r2=0.96). Pairs are coloured according to relatedness as estimated with KIN (for 1st-3rd degree); 4th-6th degree relatives annotated for pairs sharing at least two independent IBD-segments >20 cM, following Ringbauer et al[3]. Complete relatedness results shown in **Extended Data Tables 6** and **7**.
